# Normalization and variance stabilization of single-cell RNA-seq data using regularized negative binomial regression

**DOI:** 10.1101/576827

**Authors:** Christoph Hafemeister, Rahul Satija

## Abstract

Single-cell RNA-seq (scRNA-seq) data exhibits significant cell-to-cell variation due to technical factors, including the number of molecules detected in each cell, which can confound biological heterogeneity with technical effects. To address this, we present a modeling framework for the normalization and variance stabilization of molecular count data from scRNA-seq experiments. We propose that the Pearson residuals from ’regularized negative binomial regression’, where cellular sequencing depth is utilized as a covariate in a generalized linear model, successfully remove the influence of technical characteristics from downstream analyses while preserving biological heterogeneity. Importantly, we show that an unconstrained negative binomial model may overfit scRNA-seq data, and overcome this by pooling information across genes with similar abundances to obtain stable parameter estimates. Our procedure omits the need for heuristic steps including pseudocount addition or log-transformation, and improves common downstream analytical tasks such as variable gene selection, dimensional reduction, and differential expression. Our approach can be applied to any UMI-based scRNA-seq dataset and is freely available as part of the R package `sctransform`, with a direct interface to our single-cell toolkit `Seurat`.

## 1 Introduction

In the analysis and interpretation of single-cell RNA-seq (scRNA-seq) data, effective pre-processing and normalization represent key challenges. While unsupervised analysis of singlecell data has transformative potential to uncover heterogeneous cell types and states, cell-to-cell variation in technical factors can also confound these results [Vallejos et al., 2017, Stegle et al., 2015]. In particular, the observed sequencing depth (number of genes or molecules detected per cell) can vary significantly between cells, with variation in molecular counts potentially spanning an order of magnitude, even within the same cell type [The Tabula Muris Consortium, 2018]. Importantly, while the now widespread use of unique molecular identifiers (UMI) in scRNA-seq removes technical variation associated with PCR, differences in cell lysis, reverse transcription efficiency, and stochastic molecular sampling during sequencing also contribute significantly, necessitating technical correction [Hicks et al., 2017]. These same challenges apply to bulk RNA-seq workflows, but are exacerbated due to the extreme comparative sparsity of scRNA-seq data [Svensson et al., 2017].

The primary goal of single cell normalization is to remove the influence of technical effects in the underlying molecular counts, while preserving true biological variation. Specifically, we propose that a dataset which has been processed with an effective normalization workflow should have the following characteristics:

1. In general, the normalized expression level of a gene should not be correlated with the total sequencing depth of a cell. Downstream analytical tasks (dimensional reduction, differential expression) should also not be influenced by variation in sequencing depth.
2. The variance of a normalized gene (across cells) should primarily reflect biological heterogeneity, independent of gene abundance or sequencing depth. For example, genes with high variance after normalization should be differentially expressed across cell types, while housekeeping genes should exhibit low variance. Additionally, the variance of a gene should be similar when considering either deeply sequenced cells, or shallowly sequenced cells.

Given its importance, there have been a large number of diverse methods proposed for the normalization of scRNA-seq data [Bacher et al., 2017, Vallejos et al., 2015, Lun et al., 2016a, Risso et al., 2018, Lopez et al., 2018, Qiu et al., 2017]. In general, these fall into two distinct sets of approaches. The first set aims to identify ’size factors’ for individual cells, as is commonly performed for bulk RNA-seq [Love et al., 2014]. For example, BASiCS by Vallejos et al. [2015] infers cell-specific normalizing constants using spike-ins, in order to distinguish technical noise from biological cell-to-cell variability. Scran by Lun et al. [2016a] pools cells with similar library sizes and uses the summed expression values to estimate pool-based size factors, which are resolved to cell-based size factors. By performing a uniform scaling per-cell, these methods assume that the underlying RNA content is constant for all cells in the dataset, and that a single scaling factor can be applied for all genes.

Alternative normalization approaches model molecule counts using probabilistic approaches. For example, initial strategies focused on read-level (instead of UMI-level) data, and modeled the measurement of each cell as a mixture of two components: a negative binomial (NB) ‘signal’ component and a Poisson ‘dropout’ component [Kharchenko et al., 2014]. For newer measurements based on UMI, modeling strategies have focused primarily on the use of the NB distribution [Grün et al., 2014], potentially including an additional parameter to model zero-inflation (ZINB). For example, ZINB-WaVE by Risso et al. [2018] models counts as ZINB in a special variant of factor analysis. scVI and DCA also use the ZINB noise model [Lopez et al., 2018, Eraslan et al., 2019], either for normalization and dimensionality reduction in Bayesian hierarchical models, or for a denoising autoencoder. These pioneering approaches extend beyond pre-processing and normalization, but rely on the accurate estimation of per-gene error models.

In this manuscript, we present a novel statistical approach for the modeling, normalization, and variance stabilization of UMI count data for scRNA-seq. We first show that different groups of genes cannot be normalized by the same constant factor, representing an intrinsic challenge for scaling-factor based normalization schemes, regardless of how the factors themselves are calculated. We instead propose to construct a generalized linear model (GLM) for each gene with UMI counts as the response and sequencing depth as the explanatory variable. We explore potential error models for the GLM, and find that the use of unconstrained NB or ZINB models leads to overfitting of scRNA-seq data, and a significant dampening of biological variance. To address this, we find that by pooling information across genes with similar abundances, we can regularize parameter estimates and obtain reproducible error models. The residuals of our ’regularized negative binomial regression’ represent effectively normalized data values that are no longer influenced by technical characteristics, but preserve heterogeneity driven by distinct biological states. Lastly, we demonstrate that these normalized values enable downstream analyses, such as dimensionality reduction and differential expression testing, where the results are not confounded by cellular sequencing depth. Our procedure is broadly applicable for any UMI-based scRNA-seq dataset, and is freely available to users through the open-source R package `sctransform` (github.com/ChristophH/sctransform), with a direct interface to our single cell toolkit `Seurat`.

## 2 Results

### 2.1 A single scaling factor does not effectively normalize both lowly and highly expressed genes

Sequencing depth variation across single cells represents a substantial technical confounder in the analysis and interpretation of scRNA-seq data. To explore the extent of this effect and possible solutions, we examined five UMI datasets from diverse tissues, generated with both plate and droplet-based protocols. We show results on all datasets in Supplementary Note 1, but focus here on a dataset of 33,148 human peripheral blood mononuclear cells (PBMC) freely available from 10x Genomics. This dataset is characteristic of current scRNA-seq experiments; we observed a median total count of 1,891 UMI/cell, and observed 16,809 genes that were detected in at least 5 cells (Fig. 1 A,B). As expected, we observed a strong linear relationship between unnormalized expression (gene UMI count) and cellular sequencing depth. We observed nearly identical trends (and regression slopes) for genes across a wide range of abundance levels, after grouping genes into six equal-width bins based on their mean abundance (Figure 1C), demonstrating that counts from both low and high abundance genes are confounded by sequencing depth and require normalization.

**Figure 1:**
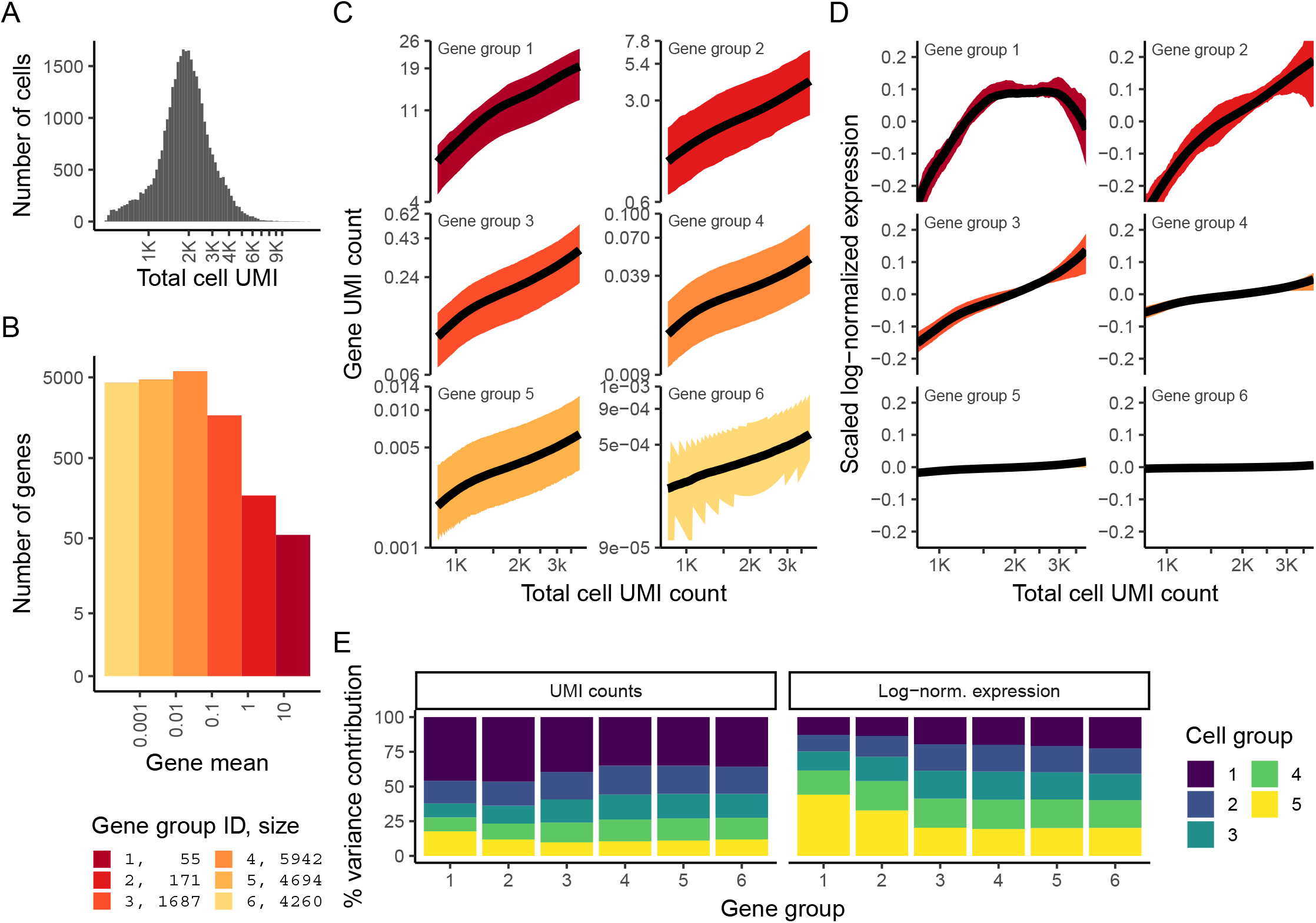
33,148 PBMC dataset from 10x genomics. **A**) Distribution of total UMI counts / cell (’sequencing depth’). **B**) We placed genes into six groups, based on their average expression in the dataset. **C**) For each gene group, we examined the average relationship between observed counts and cell sequencing depth. We fit a smooth line for each gene individually and combined results based on the groupings in (B). Black line shows mean, colored region indicates interquartile range. **D**) Same as in (C), but showing scaled log-normalized values instead of UMI counts. Values were scaled (z-scored) so that a single y-axis range could be used. **E**) Relationship between gene variance and cell sequencing depth; Cells were placed into five equal-sized groups based on total UMI counts (group 1 has the greatest depth), and we calculated the total variance of each gene group within each bin. For effectively normalized data, each cell bin should contribute 20% to the variance of each gene group.

We next tested how standard log-normalized values, the standard normalization approach in popular scRNA-seq packages such as Seurat [Satija et al., 2015, Butler et al., 2018, Stuart et al., 2018] and SCANPY [Wolf et al., 2018], compensates for this effect. As UMI counts are first scaled by the total sequencing depth (’size factors’) followed by pseudocount addition and log-transformation, we observed a weaker relationship as expected. However, we found that genes with different overall abundances exhibited distinct patterns after log-normalization, and only low/medium abundance genes in the bottom three tiers were effectively normalized (Figure 1D).

Moreover, we also found that gene variance was also confounded with sequencing depth. We quantified this phenonemon by binning cells by their overall sequencing depth, and quantifying the total variance of each gene group within each bin. For effectively normalized data, we expect uniform variance across cell groups, but we observed substantial imbalances in the analysis of log-normalized data. In particular, cells with low total UMI counts exhibited disproportionately higher variance for high-abundance genes, dampening the variance contribution from other gene groups (Figure 1E). We also tested an alternative to log-normalization (’relative counts’ normalization), where we simply divided counts by total sequencing depth. Removing the log-transformation mitigated the relationships between gene expression, gene variance, and sequencing depth, but residual effects remained in both cases (Supp. Figure 1).

These results demonstrate inherent challenges for ’size factor’-based normalization strategies. Notably, while recent normalization strategies leverage more advanced strategies to learn cell ’size factors’ [Lun et al., 2016b, Vallejos et al., 2015], the use of a single factor will introduce distinct effects on different gene sets, given their average abundance. This suggests that genes may require normalization strategies that depend on their abundance level. Indeed Bacher et al. [2017] reached similar conclusions in the normalization of non-UMI based single cell RNA-seq data. Their method, SCnorm, utilizes quantile regression to treat distinct gene groups separately, but ignores zero values which predominantly characterize droplet-based scRNA-seq. We therefore explored alternative solutions based on statistical modeling of the underlying count data.

### 2.2 Modeling single cell data with a negative binomial distribution leads to overfitting

We considered the use of generalized linear models as a statistical framework to normalize single cell data. Motivated by previous work that has demonstrated the utility of GLMs for differential expression [Finak et al., 2015], we reasoned that including sequencing depth as a GLM covariate could effectively model this technical source of variance, with the GLM residuals corresponding to normalized expression values. The choice of a GLM error model is an important consideration, and we first tested the use of a negative binomial distribution, as has been proposed for overdispersed single-cell count data [Grün et al., 2014, Risso et al., 2018], performing ’negative binomial regression’ (Methods) independently for each gene. This procedure learns three parameters for each gene, an intercept term *β*_0_ and the regression slope *β*_1_ (influence of sequencing depth), which together define the expected value, and the dispersion parameter *θ* characterizing the variance of the negative binomial errors.

We expected that we would obtain consistent parameter estimates across genes, as sequencing depth should have similar (but not identical as shown above) effects on UMI counts across different loci. To our surprise, we observed significant heterogeneity in the estimates of all three parameters, even for genes with similar average abundance (Figure 2). These differences could reflect true biological variation in the distribution of single-cell gene expression, but could also represent irreproducible variation driven by overfitting in the regression procedure. To test this, we bootstrapped the analysis by repeatedly fitting a GLM to randomized subsets of cells, and assessed the variance of parameter estimates. We found that parameter estimates were not reproducible across bootstraps (Figure 2), particularly for genes with low to moderate expression levels, suggesting that the gene-specific differences we observed were exaggerated due to overfitting.

**Figure 2:**
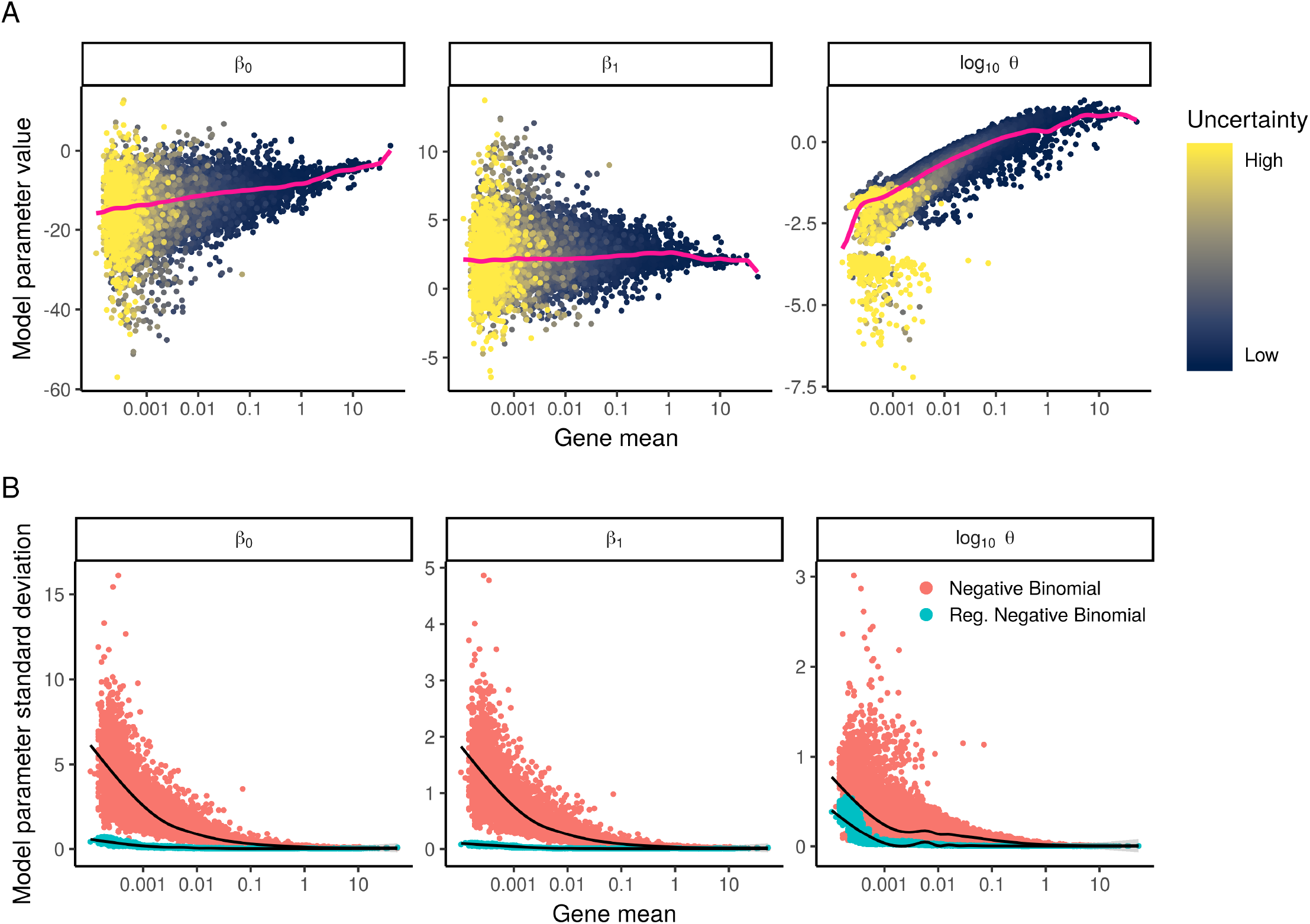
We fit NB regression models for each gene individually, and bootstrapped the process to measure uncertainty in the resulting
parameter estimates. **A**) Model parameters for 16,809 genes for the NB regression model, plotted as a function of average gene abundance. The color of each point indicates a parameter uncertainty score as determined by bootstrapping (Methods). Pink line shows the regularized parameters obtained via kernel regression. **B**) Standard deviation (σ) of NB regression model parameters across multiple bootstraps. Red points: σ for unconstrained NB model. Blue points: σ for regularized NB model, which is substantially reduced in comparison. Black trendline shows an increase in σ for low-abundance genes, highlighting the potential for overfitting in the absence of regularization.

Our observation that single cell count data can be overfit by a standard (two-parameter) NB distribution demonstrates that additional constraints may be needed to obtain robust parameter estimates. We therefore considered the possibility of constraining the model parameters through regularization, by combining information across similar genes to increase robustness and reduce sampling variation. This approach is commonly applied in learning error models for bulk RNA-seq in the context of differential expression analysis [Anders and Huber, 2010, Pimentel et al., 2017, Law et al., 2014], but to our knowledge has not been previously applied in this context for single-cell normalization.

We therefore applied kernel regression (Methods) to model the global dependence between each parameter value and average gene expression. The smoothed line (pink line in Figure 2) represents a regularized parameter estimate, that can be applied to constrain NB error models. We repeated the bootstrap procedure and found that in contrast to independent gene-level estimates, regularized parameters were consistent across repeated subsamples of the data (Supp. Figure 2), suggesting that we are robustly learning the global trends that relate intercept, slope, and dispersion to average gene expression. We note that in contrast to our approach, the use of a zero-inflated negative binomial model requires an additional (third) parameter, exacerbating the potential for over-fitting. We therefore suggest caution and careful consideration when applying unconstrained NB or ZINB models to scRNA-seq UMI count data.

### 2.3 Applying regularized negative binomial regression for single-cell normalization

Our observations above suggest a statistically-motivated, robust, and efficient process to normalize single-cell data. First, we utilize generalized linear models to fit model parameters for each gene in the transcriptome, (or a representative subset; Supp. Figure 2; Methods) using sequencing depth as a covariate. Second, we apply kernel regression to the resulting parameter estimates in order to learn regularized parameters that depend on a gene’s average expression, and are robust to sampling noise. Finally, we perform a second round of NB regression, constraining the parameters of the model to be those learned in the previous step (since the parameters are fixed, this step reduces to a simple affine transformation; Methods). We treat the residuals from this model as normalized expression levels. Positive residuals for a given gene in a given cell indicate that we observed more UMIs than expected given the gene’s average expression in the population and cellular sequencing depth, while negative residuals indicate the converse. We utilize the Pearson residuals (response residuals divided by the expected standard deviation), effectively representing a variance-stabilizing transformation (VST), where both lowly and highly expressed genes are transformed onto a common scale.

This workflow also has attractive properties for processing single cell UMI data, including:

1. We do not assume a fixed ’size’, or expected total molecular count, for any cell.
2. Our regularization procedure explicitly learns and accounts for the well-established relationship [Eling et al., 2018] between a gene’s mean abundance and variance in single cell data
3. Our VST is data driven and does not involve heuristic steps, such as a log-transformation, pseudocount addition, or z-scoring.
4. As demonstrated below, Pearson residuals are independent of sequencing depth, and can be used for variable gene selection, dimensional reduction, clustering, visualization, and differential expression.

### 2.4 Pearson residuals effectively normalize technical differences, while retaining biological variation

To evaluate our regularized NB regression model, we explored the relationship between the Pearson residuals and cellular sequencing depth. Encouragingly, we observed minimal correlation (Figure 3A,C), for genes across the full range of expression levels. In addition, gene variance was strikingly consistent across cells with different sequencing depths (Figure 3B, contrast to Figure 1E), with no evidence of expression ’dampening’ as we observed when using a cell-level size factor approaches. Taken together, these results suggest that our Pearson residuals represent effectively standardized expression values, that are not influenced by technical metrics.

**Figure 3:**
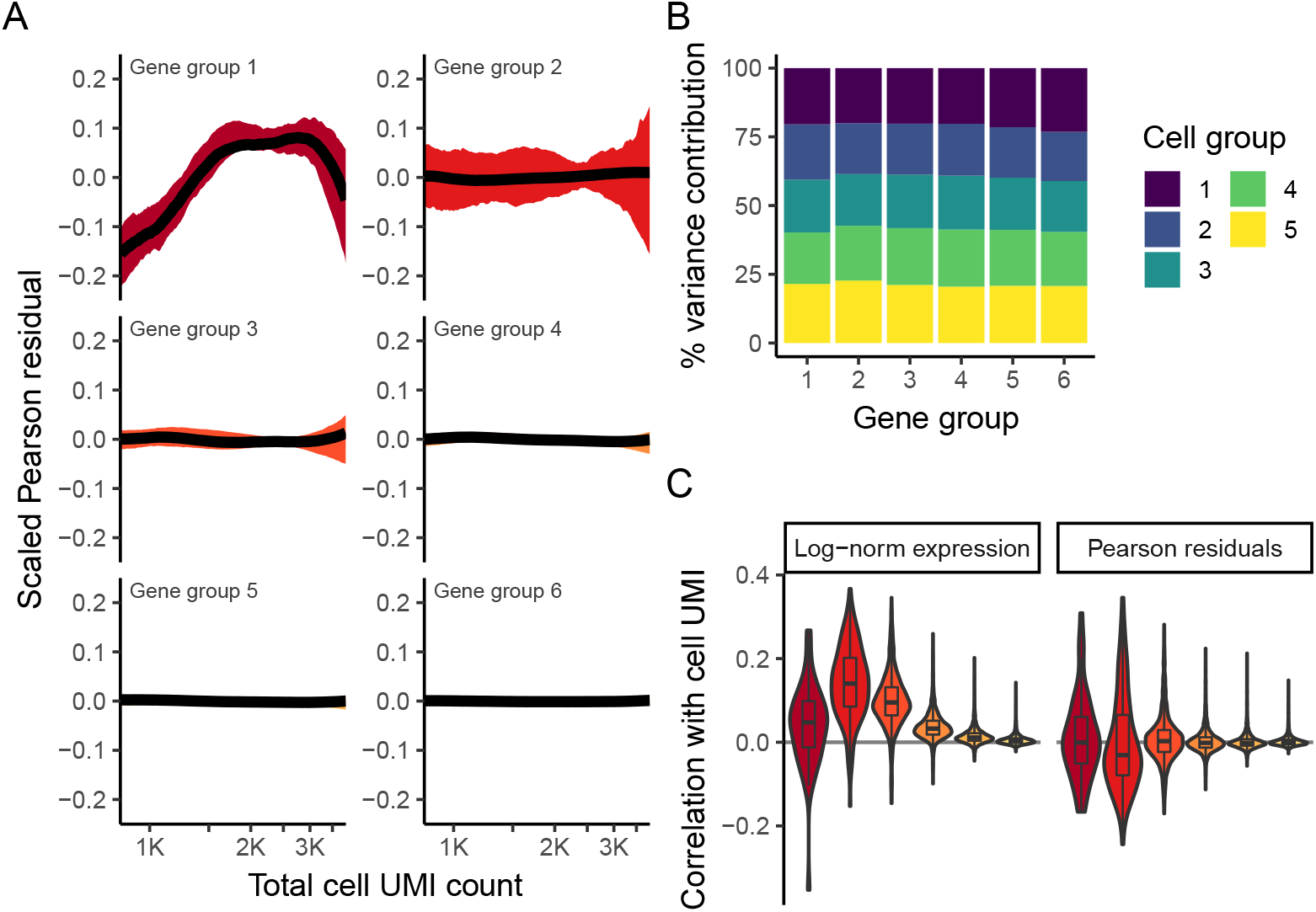
Pearson residuals from regularized NB regression represent effectively normalized scRNA-seq data. Panels **A,B** are analogous to Fig. 1D,E, but calculated using Pearson residuals. **C**) Boxplot of Pearson correlations between Pearson residuals and total cell UMI counts for each of the six gene bins. All three panels demonstrate that in contrast to log-normalized data, the level and variance of Pearson residuals is independent of sequencing depth.

Our model predicts that for genes with minimal biological heterogeneity in the data (i.e. genes whose variance is driven primarily by differences in sequencing depth), residuals should be distributed with a mean of zero and unit variance. We observe these values for the majority of genes in the dataset (Figures 4 A,B), demonstrating effective and consistent variance stabilization across a range of expression values (Figure 4C). However, we observed a set of outlier genes with substantially higher residual variance than predicted by our background model, suggesting additional biological sources of variation in addition to sampling noise. Further exploration of these genes revealed that they exclusively represent markers of known immune cell subsets (e.g. PPBP in Megakaryocytes, GNLY in NK cells, IGJ in plasma cells), demonstrating that the variance of Pearson residuals correlates with biological heterogeneity, and can be used to identify ’highly variable’ genes in single cell data. In summary, our regularized NB regression model successfully captures and removes variance driven by technical differences, while retaining biologically relevant signal.

**Figure 4:**
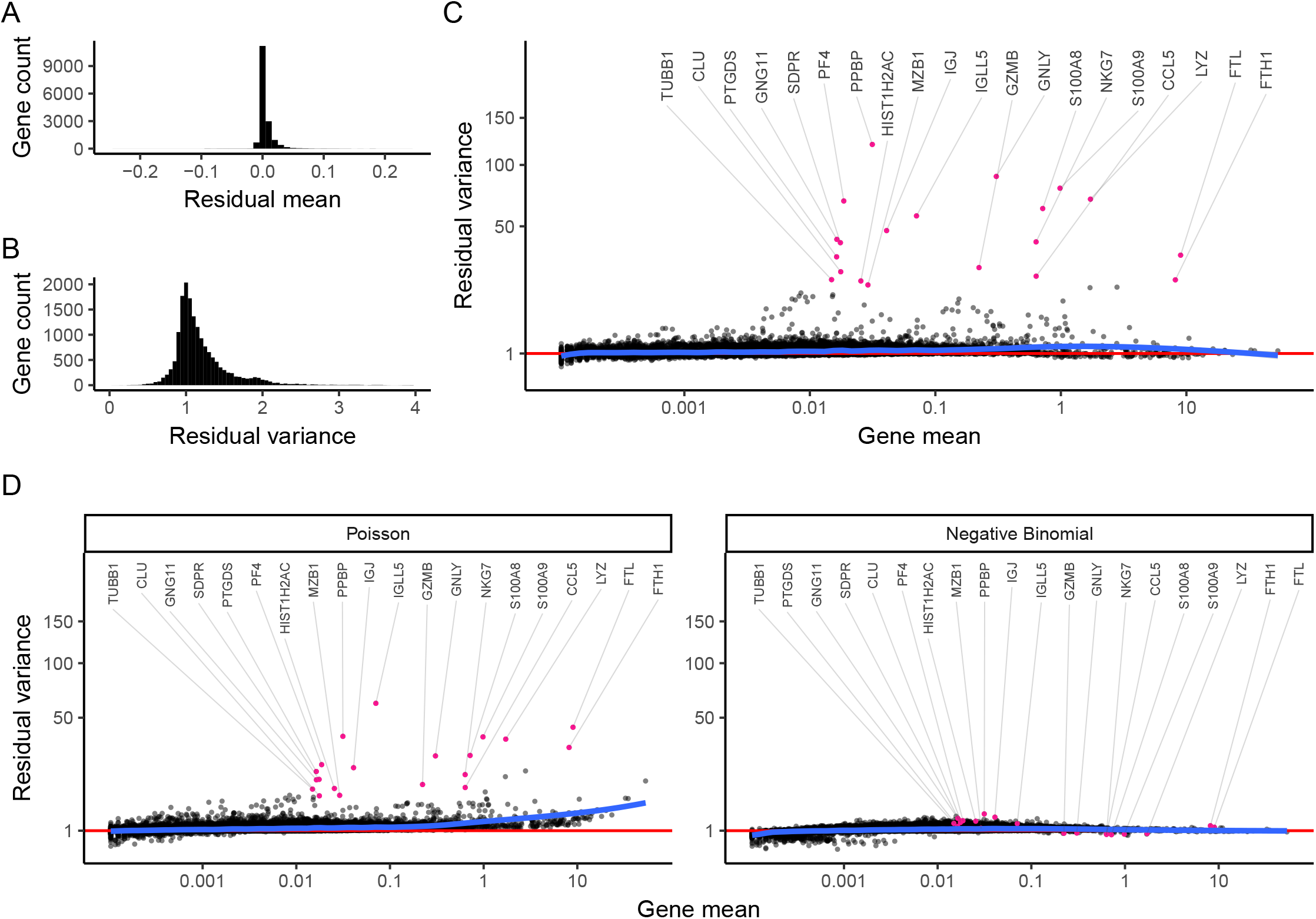
Regularized NB regression removes variation due to sequencing depth, but retains biological heterogeneity. **A**) Distribution of residual mean, across all genes, is centered at 0. **B**) Density of residual gene variance peaks at 1, as would be expected when the majority of genes do not vary across cell types. **C**) Variance of Pearson residuals is independent of gene abundance, demonstrating that the GLM has successfully captured the mean-variance relationship inherent in the data. Genes with high residual variance are exclusively cell-type markers. **D**) In contrast to a regularized NB, a Poisson error model does not fully capture the variance in highly expressed genes. An unconstrained (non-regularized) NB model overfits scRNA-seq data, attributing almost all variation to technical effects. As a result, even cell-type markers exhibit low residual variance. Mean-variance trendline shown in blue for each panel

Our previous analyses suggest that the use of a regularized NB error model is crucial to the performance of our workflow. To test this, we substituted both a Poisson and an unconstrained NB error model into our GLM, and repeated the procedure (Figure 4D). When applying standard negative binomial regression, we found that the procedure strikingly removed both technical and biological sources of variation from the data, driven by overfitting of the unconstrained distribution. A single-parameter Poisson model performed similarly to our regularized NB, but we observed that residual variances exceeded one for all moderately and highly expressed genes. This is consistent with previous observations in both bulk and single cell RNA-seq that count data is overdispersed [Risso et al., 2018, Grün et al., 2014, Love et al., 2014, Robinson et al., 2010].

In addition to global analyses, it is also instructive to explore how each model performs on characteristic genes in the dataset. In Figure 5 we show observed molecular counts for four representative loci, as a function of total cell UMI count. Background colors indicate GLM Pearson residuals values using three different error models (Poisson, NB, regularized NB), enabling us to explore how well each model fits the data. For MALAT1, a highly expressed gene that should not vary across immune cell subsets, we observe that both the unconstrained and regularized NB distributions appropriately modeled technically-driven heterogeneity in this gene, resulting in minimal residual biological variance. However, the Poisson model does not model the overdispersed counts, incorrectly suggesting significant biological heterogeneity. For S100A9 (a marker of myeloid cell types) and CD74 (expressed in antigen-presenting cells) the regularized NB and Poisson models both return bimodally distributed Pearson residuals, consistent with a mixture of myeloid and lymphoid cell types present in blood, while the unconstrained NB collapses this biological heterogeneity via overfitting. We observe similar results for the Megakaryocyte (Mk) marker PPBP, but note that both non-regularized models actually fit a negative slope relating total sequencing depth to gene molecule counts. This is because Mk cells have very little RNA content, and therefore exhibit lower UMI counts compared to other cell types, even independent of stochastic sampling. However, it is nonsensical to suggest that deeply sequenced Mk cells should contain less PPBP molecules than shallowly sequenced Mk cells, and indeed, regularization of the slope parameter overcomes this problem.

**Figure 5:**
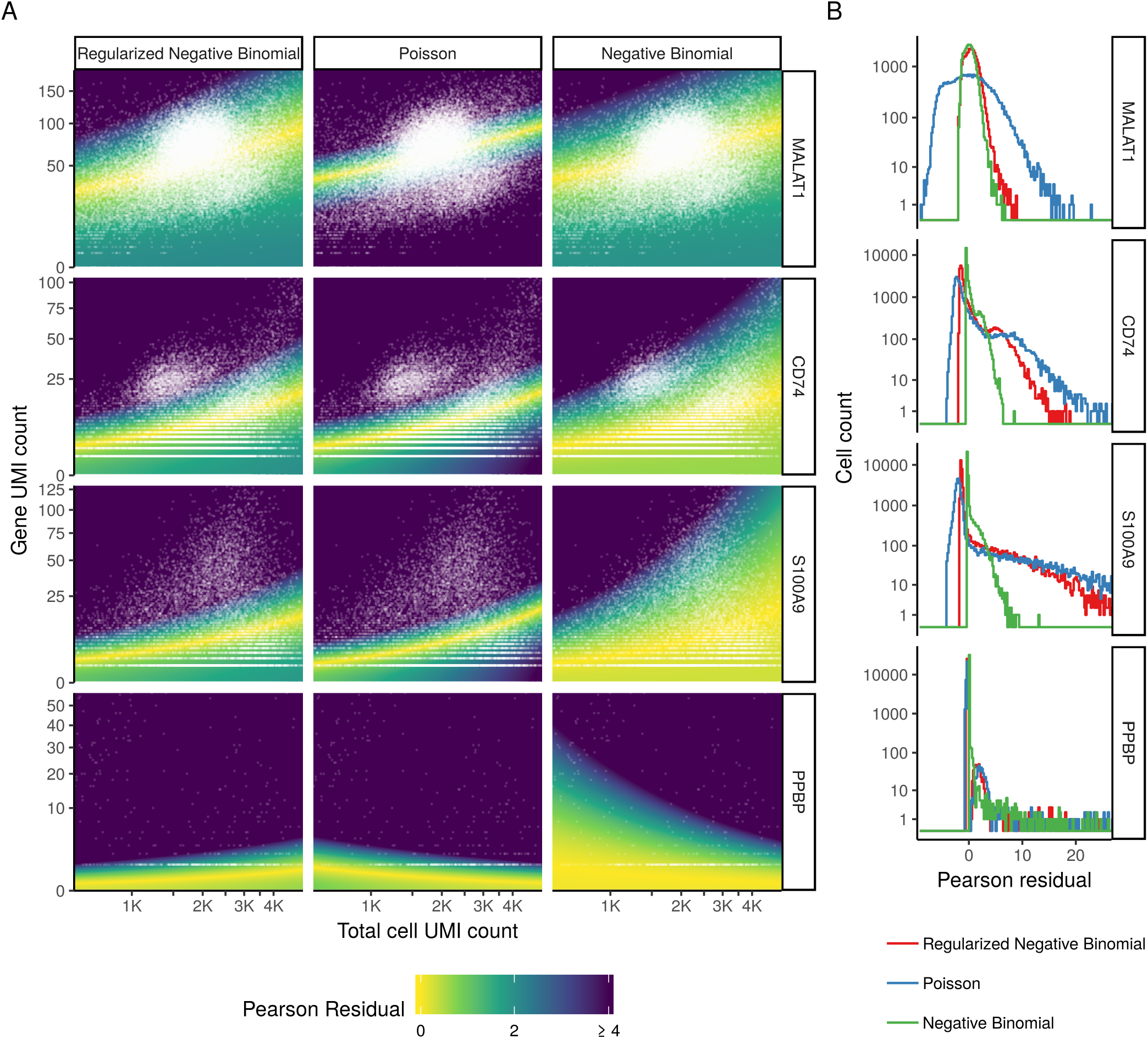
The regularized NB model is an attractive middle ground between two extremes. **A**) For four genes, we show the relationship between cell sequencing depth and molecular counts. White points show the observed data. Background color represents the Pearson residual magnitude under three error models. For MALAT1 (does not vary across cell types) the Poisson error model does not account for overdispersion, and incorrectly infers significant residual variation (biological heterogeneity). For S100A9 (a CD14+ Monocyte marker) and CD74 (expressed in antigen-presenting cells) the non-regularized NB model overfits the data, and collapses biological heterogeneity. For PPBP (a Megakaryocyte marker) both non-regularized models wrongly fit a negative slope. **B**) Boxplot of Pearson residuals for models shown in A. X-axis range shown is limited to [-8, 25] for visual clarity.

Taken together, our results demonstrate that the regularized negative binomial represents an attractive middle ground between two extremes. By allowing for overdispersion, the model can correctly account for the variance in count data observed in single cell assays. However, by placing data-driven constraints on the slope, intercept, and dispersion of NB regression, we substantially alleviate the problem of overfitting, and ensure that biological variation is retained after normalization. We observed identical results when considering any of the five UMI datasets we tested, including both plate and droplet-based protocols (Supplementary Note 1), demonstrating that our procedure can apply widely to any UMI-based scRNA-seq experiment.

### 2.5 Downstream analytical tasks are not biased by sequencing depth

Our procedure is motivated by the desire to standardize expression counts so that differences in cellular sequencing depth do not influence downstream analytical tasks. To test our performance towards this goal, we performed dimensionality reduction and differential expression tests on Pearson residuals after regularized NB regression. For comparison, we used log-normalized data. We first applied PCA followed by UMAP embedding (Methods) to the full PBMC dataset, using normalized values (Pearson residuals, or log-normalized data) for all genes in the transcriptome as input to PCA, and then visualized the total number of molecules per-cell on the UMAP embedding. Both normalization schemes reveal significant biological heterogeneity in PBMC (Figure 6A), consistent with the expected major and minor human immune cell subsets. However, the low-dimensional representation of log-normalized data was confounded by cellular sequencing depth, as both the PCA and UMAP embeddings exhibited strong correlations with this technical metric. These correlations are strikingly reduced for Pearson residuals (Figure 6A). We note that we do not expect complete independence of biological and technical effects, as distinct cell subsets will likely vary in size and RNA content. However, even when limiting our analyses within individual cell types, we found that cell sequencing depth explained substantially reduced variation in Pearson residuals compared to log-normalized data (Figure 6B), consistent with our earlier observations (Figure 3).

**Figure 6:**
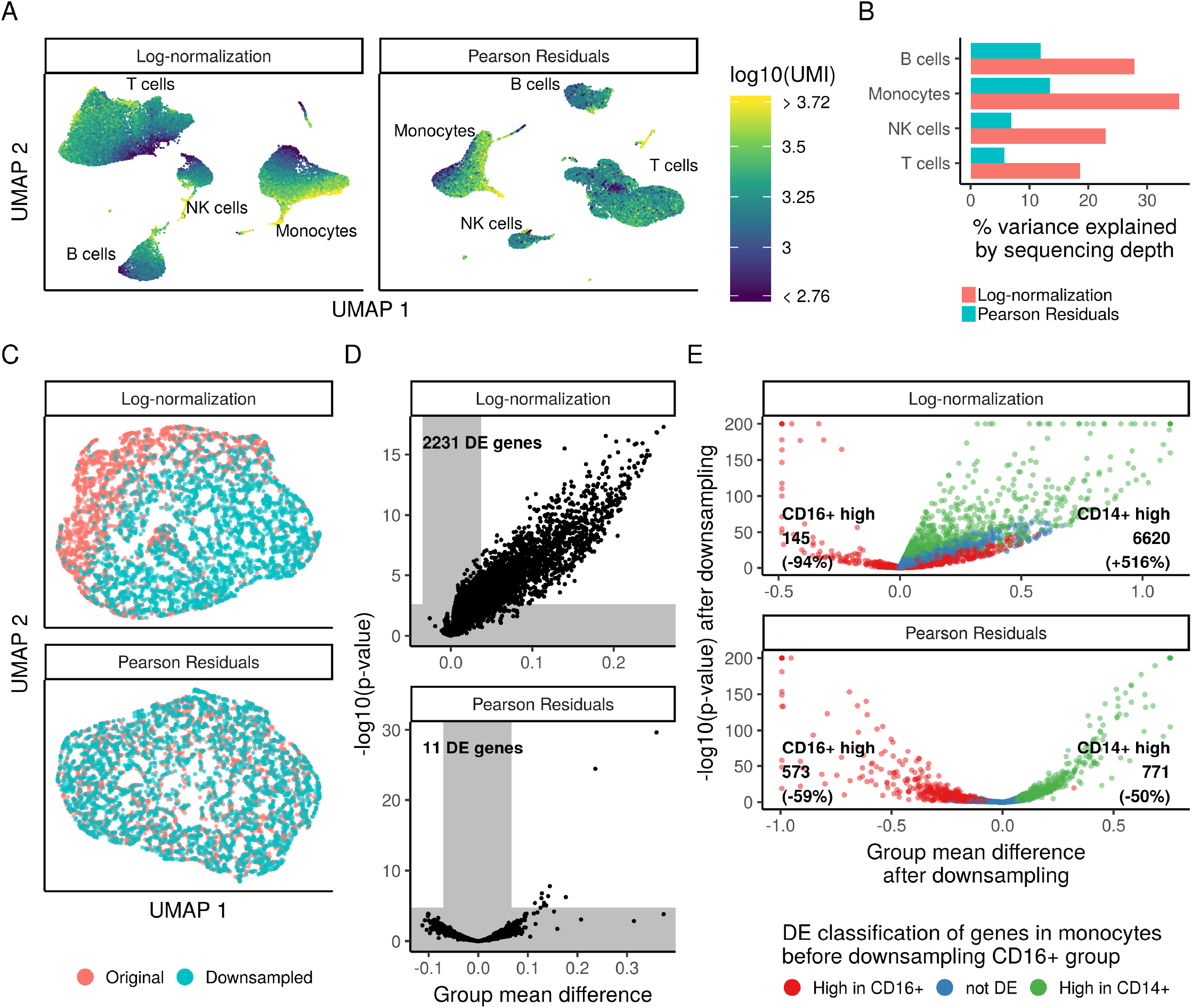
Downstream analyses of Pearson residuals are unaffected by differences in sequencing depth. **A**) UMAP embedding of the 33,148 cell PBMC dataset using either log-normalization or Pearson residuals. Both normalization schemes lead to similar results with respect to the major and minor cell populations in the dataset. However, in analyses of log-normalized data, cells within a cluster are ordered along a gradient that is correlated with sequencing depth. **B**) Within the four major cell types, the percent of variance explained by sequencing depth under both normalization schemes. **C**) UMAP embedding of two groups of biologically identical CD14+ monocytes, where one group was randomly downsampled to 50% depth. **D**) Results of differential expression (DE) test between the two groups shown in C. Gray areas indicate expected group mean difference by chance and a false discovery rate cutoff of 1%. **E**) Results of DE test between CD14+ and CD16+ monocytes, before and after randomly downsampling the CD16+ cells to 20% depth.

Imperfect normalization can also confound differential expression (DE) tests for scRNA-seq, particularly if global differences in normalization create DE false positives for many genes. To demonstrate the scope of this problem, and test its potential resolution with Pearson residuals, we took CD14+ monocytes (5,551 cell subset of the 33K PBMC data) and randomly divided them into two groups. In one of the groups (50% of the cells), we randomly subsampled UMIs so that each cell expressed only 50% of its total UMI counts. Therefore, the two groups of monocytes are biologically equivalent, and differ only in their technical sequencing depth, and we should ideally detect no differentially expressed genes between them. However, when performing DE on log-normalized data (t-test with significance thresholds determined by random sampling, see Methods), we detected more than 2,000 DE genes (FDR threshold 0.01), due to global shifts arising from improper normalization (Figure 6C,D). When performing DE on Pearson residuals, we identified only 11 genes. While these 11 represent false positives, they are each highly expressed genes for which it is difficult to obtain a good fit during the regularization process as there are few genes with similar mean values (Figure 3A top left).

We also tested a second scenario where true DE genes could be masked by sequencing depth differences. We compared two distinct populations, CD14+ and CD16+ monocytes (5,551 and 1,475 cells), before and after randomly downsampling the CD16+ group to 20% sequencing depth. We would expect the set of DE genes to be nearly identical in the two analyses, though we expect a decrease in sensitivity after downsampling. However, when using log-normalized data, we observed dramatic changes in the set of DE genes - with some CD14+-enriched markers even incorrectly appearing as CD16+-enriched markers after downsampling. When performing DE on Pearson residuals, the results of the two analyses were highly concordant, albeit with reduced statistical power after downsampling (Figure 6E). Therefore, Pearson residuals resulting from regularized NB regression effectively mitigate depth-dependent differences in dimensionality reduction and differential expression, which are key downstream steps in single cell analytical workflows.

## 3 Discussion

Here, we present a statistical approach for the normalization and variance stabilization of single cell UMI datasets. In contrast to commonly applied normalization strategies, our workflow omits the use of linear size/scaling factors, and focuses instead on the construction of a GLM relating cellular sequencing depth to gene molecule counts. We calculate the Pearson residuals of this model, representing a variance-stabilization transformation that removes the inherent dependence between a gene’s average expression and cell-to-cell variation. We demonstrate that our Pearson residuals represent normalized scRNA-seq values, that can be utilized for diverse downstream tasks including variable gene selection, dimensional reduction, and differential expression. In each case, our procedure effectively removes the influence of technical variation, without dampening biological heterogeneity.

When exploring error models for the GLM, our analyses revealed that an unconstrained negative binomial model tends to overfit single cell RNA-seq data, particularly for genes with low/medium abundance. We demonstrate that a regularization step, a commmon step in bulk RNA-seq analysis [Robinson et al., 2010, McCarthy et al., 2012] where parameter estimates are pooled across genes with similar mean abundance, can effectively overcome this challenge and yield reproducible models. Importantly, many statistical and deep-learning methods designed for single cell RNA-seq data often utilize a negative binomial (or zero-inflated negative binomial) error model [Lopez et al., 2018, Eraslan et al., 2019]. Our results suggest that these and future methods could benefit by substituting a regularized model, and that including an additional parameter for zero-inflation could exacerbate the risk of overfitting. More generally, our work indicates that a regularized negative binomial is an appropriate distribution to model UMI count data from a ’homogeneous’ single cell population.

To facilitate users applying these methods to their own datasets, our approach is freely available as an open-source R package `sctransform` (github.com/ChristophH/sctransform), with an accompanying interface to our single cell R toolkit Seurat [Satija et al., 2015, Butler et al., 2018, Stuart et al., 2018]. In a single command, and without any requirement to set user-defined parameters, `sctransform` performs normalization, variance stabilization, and feature selection based on a UMI-based gene expression matrix. We demonstrate the ease-of-use for `sctransform` in a short vignette analyzing a 2,700 PBMC dataset produced by 10x Genomics in Supplementary Note 2. In this example, `sctransform` reveals significant additional biological substructure in NK, T, B, and monocyte populations that cannot be observed in the standard `Seurat` workflow, which is based on log-normalization (Supplementary Note 2).

As our workflow leverages all genes (or a random subset) for the initial regularization, we make an implicit assumption that the majority of genes in the dataset do not exhibit significant biological variation. This is analogous to similar assumptions made for bulk RNA-seq normalization and DE (i.e., that the majority of genes are not differentially expressed across conditions) [Robinson et al., 2010]. While this assumption may be overly simplistic when performing scRNA-seq on a highly heterogeneous sample, we did not observe adverse affects when applying our model to human PBMC data, or any of the other four datasets we examined. In principle, an initial pre-clustering step (as proposed in [Lun et al., 2016a]) could alleviate this concern, as the biological heterogeneity would be significantly reduced in each group.

Finally, while we focus here on modeling technical variation due to differences in cellular sequencing depth, we note that our approach can be easily extended to model alternative ’nuisance’ parameters, including cell cycle [Buettner et al., 2015], mitochondrial percentage, or experimental batch, simply by adding an additional covariates to the model. Indeed, we observed that a modified GLM including a batch indicator variable was sufficient to correct for technical differences arising across from two profiled batches of murine bipolar cells [Shekhar et al., 2016], though successful application requires all cell types to share a similar batch effect (Supp. Figure 3). In the future, we anticipate that similar efforts can be used to model diverse single-cell data types, including single-cell protein [Stoeckius et al., 2017], chromatin [Buenrostro et al., 2015], and spatial [Wang et al., 2018] data.

## 4 Methods

### 4.1 Regularized negative binomial regression

We explicitly model the UMI counts for a given gene using a generalized linear model. Specifically we use the sum of all molecules assigned to a cell as a proxy for sequencing depth, and use this cell attribute in a regression model with negative binomial (NB) error distribution and log link function. Thus, for a given gene *i*, we have

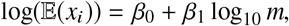

where *x_i_* is the vector of UMI counts assigned to gene *i*, and m is the vector of molecules assigned to the cells, i.e. 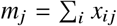. The solution to this regression is a set of parameters: the intercept *β*_0_, the slope *β*_1_. The dispersion parameter *θ* of the underlying NB distribution is also unknown and needs to be estimated from the data. Here we use the NB parameterization with mean *μ* and variance given as 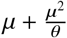.

We use a regression model for the UMI counts to correct for sequencing depth differences between cells, and to standardize the data. However, modeling each gene separately results in overfitting, particularly for low-abundane genes that are detected in only a minor subset of cells, and are modeled with a high variance. We consider this an overestimation of the true variance, as this is drive by cell type heterogeneity in the sample, and not due to cell-to-cell variability with respect to the independent variable, log_10_ *m*. To avoid this overfitting, we regularize all model parameters, including the NB dispersion parameter *θ*, by sharing information across genes.

The procedure we developed has three steps. In the first step, we fit independent regression models per gene. In the second step, we exploit the relationship of model parameter values and gene mean to learn global trends in the data. We capture these trends using a kernel regression estimate (ksmooth function in R) with normal kernel and large bandwidth (3 times the size as suggested by R function bw.SJ). We perform independent regularizations for all parameters (Fig 2). In the third step, we use the regularized regression parameters to define an affine function that transforms that converts UMI counts into Pearson residuals:

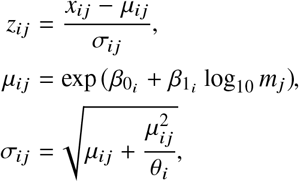

where *z_ij_* is the Pearson residual of gene *i* in cell *j*, *x_ij_* is the observed UMI count of gene *i* in cell *j*, *μ_ij_* is the expected UMI count of gene *i* in cell *j* in the regularized NB regression model, and *σ_ij_* is the expected standard deviation of gene *i* in cell *j* in the regularized NB regression model. Here *β*_0_*i*__, *β*_1_*i*__, and *θ_i_* are the linear model parameters after regularization. To reduce the impact of extreme outliers, we clip the residuals to a maximum value of 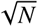, where *N* is the total number of cells.

### 4.2 Geometric mean for genes

Our regularization approach aims to pool information across genes with similar average expression. To avoid the influence of outlier cells and respect the exponential nature of the count distributions, we consistently use the geometric mean. References instances that refer to average abundance or gene mean in this work are based on the following definition of mean:

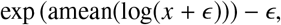

with *x* being the vector of UMI counts of the gene, amean being the arithmetic mean, and *ϵ* being a small fixed value to avoid log(0). After trying several values for *ϵ* in the range 0.0001 to 1, and not observing significant differences in our results, we set *ϵ* = 1.

### 4.3 Speed Considerations

`sctransform` has been optimized to run efficiently on large scRNA-seq datasets on standard computational infrastructure. For example, processing of a 3,000 cell dataset takes 30 seconds on a standard laptop (the 33,148 cell dataset utilized in this manuscript takes 6 minutes).

The most time consuming step of our procedure is the initial GLM-fitting, prior to regularization. Here, we fit *K* linear regression models with NB error models, where *K* is the total number of genes in the dataset. However, since the results of the first step are only used to learn regularized parameter estimates (ie. the overall relationship of model parameter values and gene mean), we tested the possibility of performing this step on a random subset of genes in lieu of the full transcriptome. When selecting a subset of genes to speed up the first step, we do not select genes at random, i.e. with a uniform sampling probability, as that would not evenly cover the range of gene means. Instead, we set the probability of selecting a gene *i* to 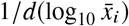, where *d* is the density estimate of all *log*_10_-transformed gene means and 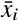 is the mean of UMI counts of gene *i*.

For different numbers of genes (ranging from 4,000 to 50), we drew 13 random samples to be used in the initial step of parameter estimation. We then proceeded to generate regularized models (for all genes based on parameters learned from a gene subset), and compared the results to the case where all genes were used in the initial estimation step as well. We employed a few metrics to compare the partial analysis to the full analysis: the correlation of gene-residuals, the ranking of genes based on residual variation (most highly-variable genes), and the CV of sum of squared residuals across random samples (model stability). For all metrics, we observed that using as few as 200 genes in the initial estimation closely recapitulated the full results, while using 2,000 genes gave rise to virtually identical estimates (Supp. Figure 2). We therefore use 2,000 genes in the initial GLM-fitting step.

Additionally, we explored three methods to estimate the model parameters in the initial step. We list them here in increasing order of computational complexity.

1. Assume a Poisson error distribution to estimate *β* coefficients. Then, given the estimated mean vector, estimate the NB *θ* parameter using maximum likelihood.
2. Same as above, followed by a re-estimation of *β* coefficients using a NB error model with the previously estimated *θ*.
3. Fit a NB GLM estimating both the *β* and *θ* coefficients using an alternating iteration process.

While the estimated model parameters can vary slightly these methods, the resulting Pearson residuals are extremely similar. For example, when applying the three procedures to the 10x PBMC dataset, all pairwise gene correlations between the three methods are greater than 0.99, though the alternating iteration process is four-fold more computationally demanding. We therefore proceeded with the first method.

### 4.4 Trends in the data before and after normalization

We grouped genes into six bins based on log_10_-transformed mean UMI count, using bins of equal width. To show the overall trends in the data, for every gene we fit the expression (UMI counts, scaled log-normalized expression, scaled Pearson residuals) as a function of log_10_-transformed mean UMI count using kernel regression (ksmooth function) with normal kernel and large bandwidth (20 times the size as suggested by R function bw.SJ). For visualization we only used the central 90% of cells based on total UMI. For every gene group we show the expression range after smoothing from first to third quartile at 200 equidistant cell UMI values.

### 4.5 Model parameter stability

To assess model parameter stability we bootstrapped the parameter estimation and sampled from all cells with replacement 13 times. For a given gene and parameter combination we derived an uncertainty score as follows. We used the standard deviation of parameter estimates across 13 bootstraps divided by the standard deviation of the bootstrap-mean value across all genes. Values greater or equal to one indicate high uncertainty, while values less or equal to 0.01 indicate low uncertainty.

### 4.6 Variance contribution analysis

To evaluate whether gene variance is dependent on sequencing depth, we determined the contribution of different cell groups to the overall variance of our six previously determined gene sets. For this we placed all cells into five equal-sized groups based on total UMI counts (group 1 has the greatest depth, group 5 the lowest). We center each gene and square the values to obtain the squared deviation from the mean. The variance contribution of a cell group is then the sum of the values in those cells divided by the sum across all cells.

### 4.7 Density maps for Pearson residuals

To illustrate different models (regularized NB, Poisson, nonregularized NB) for four example genes, we show Pearson residuals on 256×256 grids in form of heatmaps. X and Y axis ranges were chosen to represent the central 98% of cells and central 99.8% of UMI counts. Heatmap colors show the magnitude (absolute value) of Pearson residuals, clipped to a maximum value of 4.

### 4.8 Dimensionality reduction

For both log-normalized data and Pearson residuals we performed dimensionality reduction as follows. We centered and scaled all 16K genes, clipped all values to the interval [-10, 10] and performed a truncated principal components analysis as provided by the irlba R package. In both cases we kept the first 25 PCs based on eigenvalue drop-off. For 2D visualization the PC embeddings were passed into UMAP [McInnes and Healy,2018, McInnes et al., 2018] with default parameters.

### 4.9 Differential expression testing

Differential expression testing was done using independent t-tests per gene for all genes detected in at least 5 cells in at least one of the two groups being compared. P-values were adjusted for multiple comparisons using the Benjamini & Hochberg method (FDR). Input to the test was either log-normalized (log(10,000UMI_gene_/UMI_cell_ + 1)) expression or Pearson residuals after regularized NB regression. A random background distribution of mean differences was generated by randomly choosing 1,000 genes and permuting the group labels. Significance thresholds for the difference of means were derived from the background distribution by taking the 0.5th and 99.5th percentile. Finally, we called genes differentially expressed if the FDR was below 0.01 and the difference of means exceeded the threshold for significance.

### 4.10 Model extensions - additional nuisance parameters

For the results shown in this manuscript, we have used the log-transformed total number of UMI assigned to each cell as the dependent variable to model gene-level UMI counts. However, other variables may also be suitable as long as they capture the sampling depth associated with each cell. While our approach is flexible, we use molecular depth as the default covariate in all of our analyses here.

Additionally, the model can be flexibly extended to include additional covariates representing nuisance sources of variation, including cell-cycle state, mitochondrial percentage, or experimental batch. In these cases (unlike with sequencing depth), no regularization can be performed for parameters involving these variables, as genes with similar abundances cannot be assumed to (for example) be expressed in a similar pattern across the cell cycle. In these cases, we first learn regularized models using only the sequencing depth covariate, as described above. We next perform a second round of NB regression, including both the depth covariate and additional nuisance parameters as model predictors. In this round, the depth-dependent parameters are fixed to their previously regularized values, while the additional parameters are unconstrained and fit during the regression. The Pearson residuals of this second round of regression represent normalized data.

As a proof-of-concept, we illustrate a potential model extension by including a batch indicator variable when analyzing a dataset of 26,439 murine bipolar cells produced by two experimental batches [Shekhar et al., 2016], considering all bipolar cells and Müller glia. After running `sctransform`, either with the inclusion or exclusion of the batch covariate, we performed PCA on all genes, and used the first 20 dimensions to compute a UMAP embedding (Supp. Figure 3). We include this example as a demonstration for how additional nuisance parameters can be included in the GLM framework, but note that when cell-type specific batch effects are present, or there is a shift in the percentage of cell types across experiments, non-linear batch effect correction strategies are needed [Stuart et al., 2018]

## Supporting information

Supplementary Note 1: Application of sctransform to additional UMI datasets

Supplementary Note 2 : Using sctransform in Seurat

## Data and Software Availability

The dataset used in the main text is ‘33k PBMCs from a Healthy Donor, v1 Chemistry’ from 10x Genomics. Additional datasets used in the study are listed in Supplementary Note 1, along with GEO accession numbers and download links.

Software implementing our approach is freely available as an open-source R package `sctransform` (github.com/ChristophH/sctransform).

## Author contributions

C.H. and R.S. conceived of the method, derived the models, performed the experiments and analysed the data. Both authors discussed the results and wrote the final manuscript.

## Competing interests

The authors declare no competing interests.

## Grant information

This work was supported by an NIH New Innovator Award (1DP2HG009623-01; R.S.), NIH Human Biomolecular Atlas Program Award (1OT2OD026673-01; R.S.), Chan Zuckerberg Award (HCA2-A-1708-02755; R.S.), and a Deutsche Forschungsgemeinschaft (DFG) Research Fellowship (328558384; C.H.).

## Acknowledgements

The authors would like to thank Joshua Batson (Chan Zuckerberg Biohub), Valentine Svennson (Caltech), Ken Harris (UCL), David Knowles (NYGC), and members of the Satija Lab for thoughtful discussions related to this work.

## Supplementary Figures

**Supp. Figure 1:**
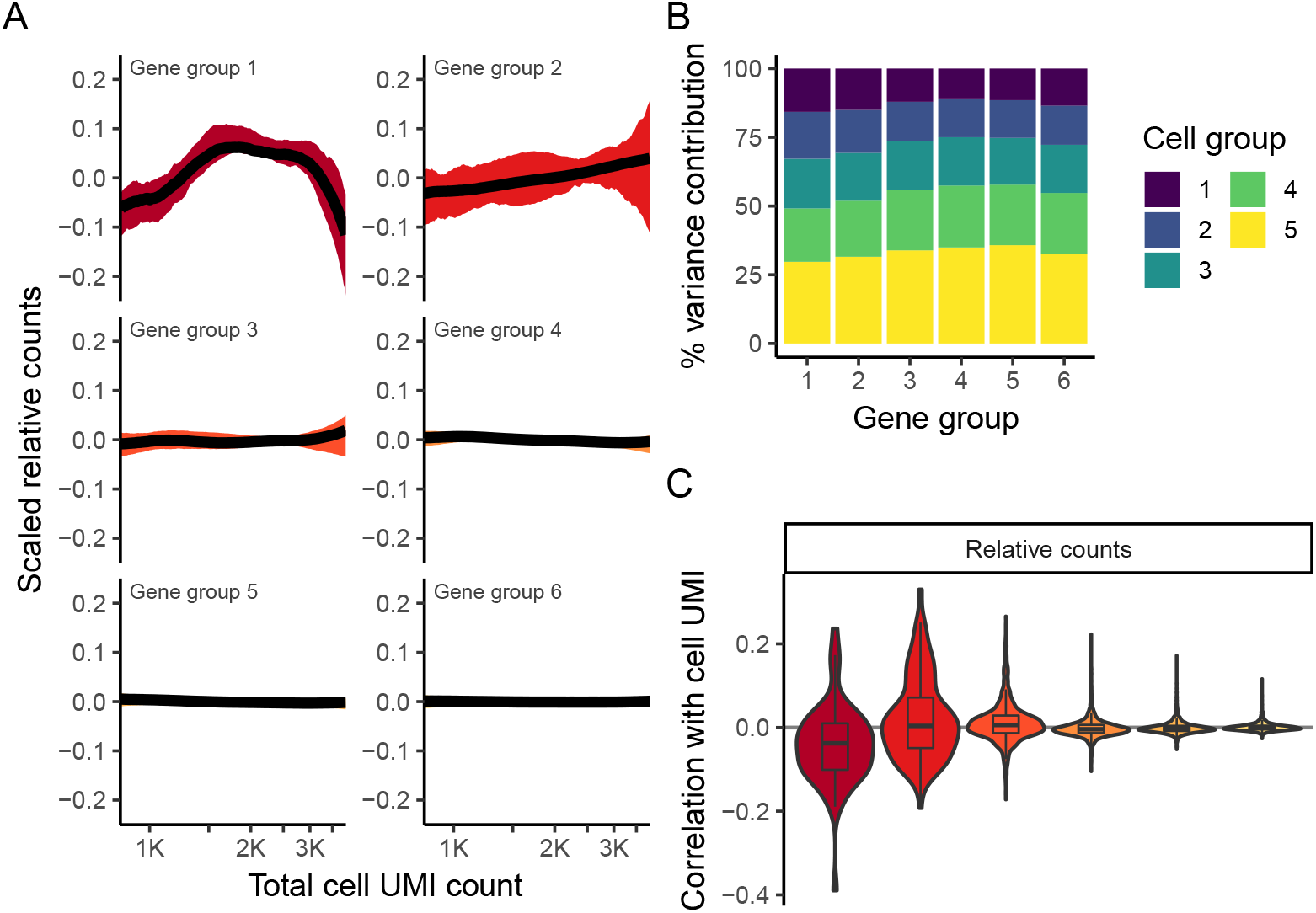
’Relative counts’ normalization of 33K PBMC data set. **A,B**) Visualizations are analogous to Fig. 1D,E, but calculated using ’relative counts’ normalization. **C**) Boxplot of Pearson correlations between ’relative counts’ and total cell UMI counts for each of the six gene bins.

**Supp. Figure 2:**
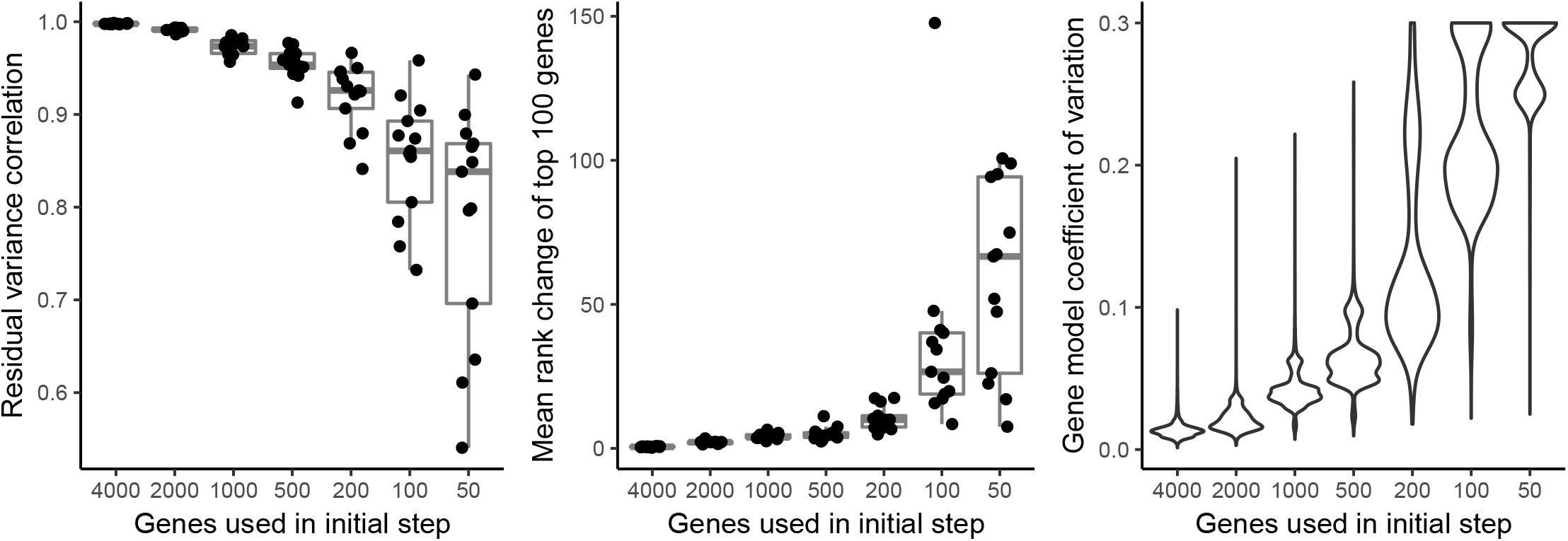
A representative set of 2,000 genes is sufficient for learning regularized models. **A,B**) Comparing models learned using only a subset of genes and models learned using all 16,809 genes. **A**) Pearson correlation of gene residuals **B**) Mean rank change of top 100 variable genes as determined by residual variance. **C**) Coefficient of variation of sum of squared residuals across multiple samples; All panels show results of 13 random samples per gene subset size

**Supp. Figure 3:**
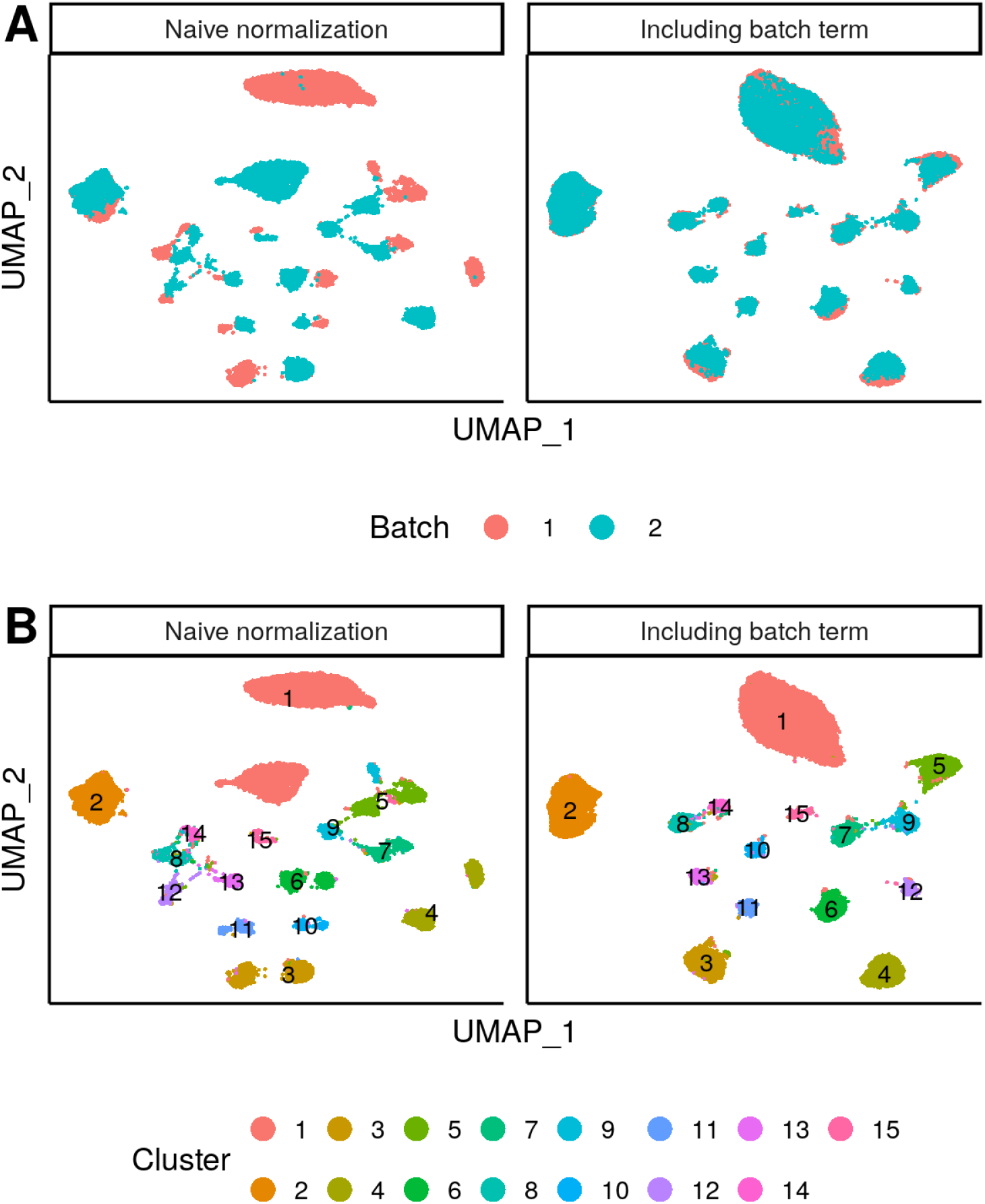
Batch-correction during normalization. **A**) UMAP embedding of the bipolar cell dataset before and after including a batch term during normalization. When applying `sctransform` without the batch indicator variable (i.e. batch-naive normalization), we see clear separation per batch, but when including a batch term in the regression model used for normalization, the batches align. **B**) Same as above, but colors indicate clusters of the original study. We include this example as a demonstration for how additional nuisance parameters can be included in the GLM framework, but note that when cell-type specific batch effects or present, or there is a shift in the percentage of cell types across experiments, non-linear batch effect correction strategies are needed. Stuart et al. [2018]

