## Supplementary Note 1: Application of sctransform to additional UMI datasets for "Normalization and variance stabilization of single-cell RNA-seq data using regularized negative binomial regression"

The following figures show results of our analysis and normalization method for four additional datasets:

1. Human pancreas (CEL-seq2) GSE85241 [Muraro et al., 2016]
2. 10X Genomics 10k PBMCs from a Healthy Donor (v3 chemistry)
3. 10X Genomics 10k Brain Cells from an E18 Mouse (v3 chemistry)
4. 10X Genomics 10k Heart Cells from an E18 mouse (v3 chemistry)

10X Genomics datasets can be found at `https://support.10xgenomics.com/single-cell-gene-expression/datasets/`

### Figure legends

#### First figure – Overview of datasets and bias in log-normalized expression

Top Distribution of total UMI counts / cell ('sequencing depth').

Middle We placed genes into six groups, based on their average expression in the dataset.

Bottom For each gene group, we examined the average relationship between log-normalized expression and cell sequencing depth. We fit a smooth line for each gene individually and combined results based on the groupings. Black line shows mean, colored region indicates interquartile range. Values were scaled (z-scored) so that a single y-axis range could be used.

#### Remaining figures – Details of normalization and variance stabilization

A) Model parameters for genes detected in at least 0.1% of the cells for the NB regression model, plotted as a function of average gene abundance. The color of each point indicates a parameter uncertainty score as determined by bootstrapping (Methods). Pink line shows the regularized parameters obtained via kernel regression.

B) Standard deviation ($\sigma$) of NB regression model parameters across multiple bootstraps. Red points: $\sigma$ for unconstrained NB model. Blue points: $\sigma$ for regularized NB model, which is substantially reduced in comparison. Black trendline shows an increase in $\sigma$ for low-abundance genes, highlighting the potential for overfitting in the absence of regularization.

C) Distribution of residual mean, across all genes, is centered at 0.

D) Density of residual gene variance peaks at 1, as would be expected when the majority of genes do not vary across cell types.

E) Variance of Pearson residuals is independent of gene abundance, demonstrating that the GLM has corrected for the mean-variance relationship inherent in the data. Genes with high residual variance tend to be cell-type markers. In contrast to a regularized NB, a Poisson error model does not fully capture the variance in highly expressed genes. An unconstrained (non-regularized) NB model overfits scRNA-seq data, attributing almost all variation to technical effects. As a result, even cell-type markers exhibit low residual variance. Mean-variance trendline shown in blue for each panel.

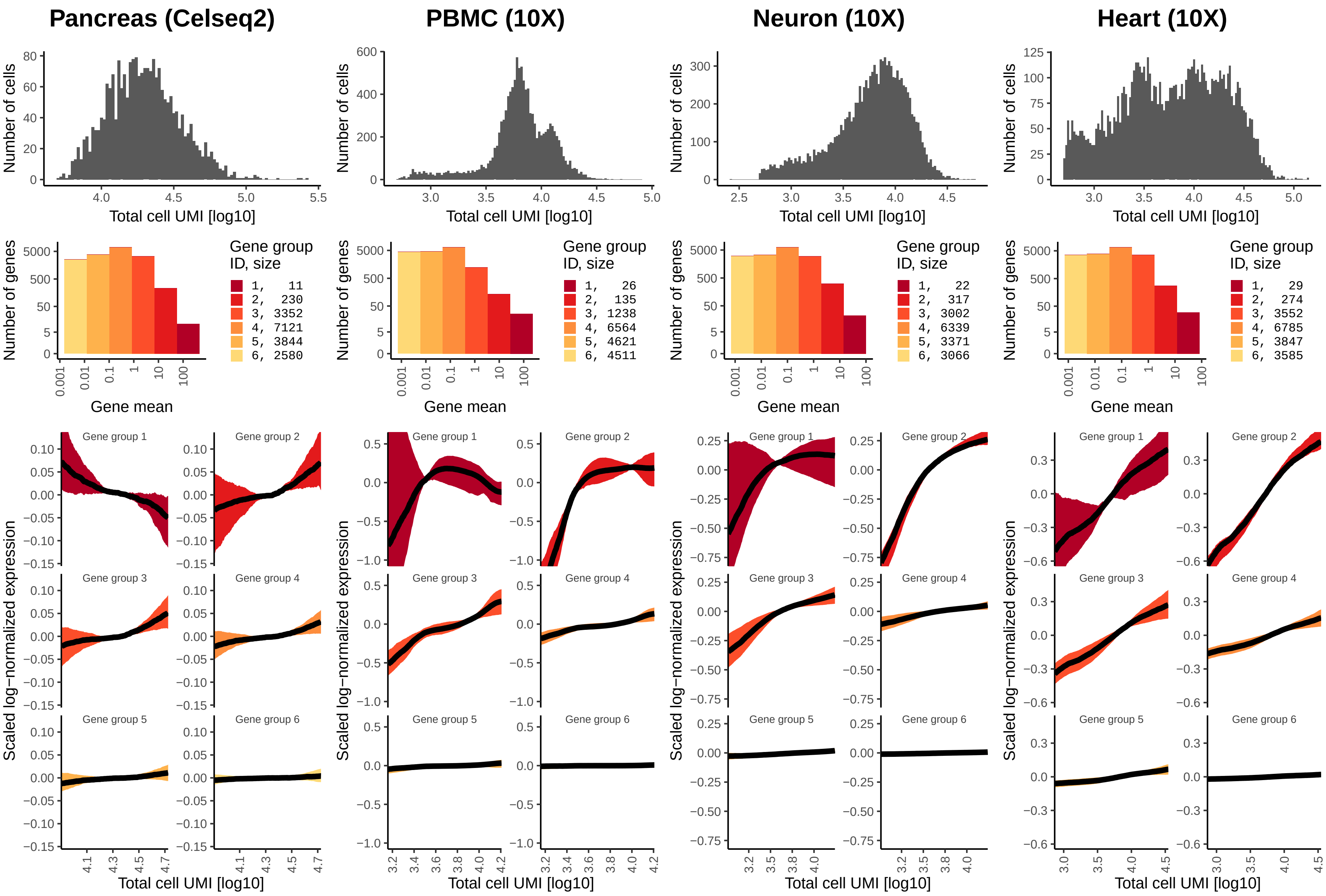
Pancreas (Celseq2)
PBMC (10X)
Neuron (10X)
Heart (10X)
Number of cells
Total cell UMI [log10]
Number of genes
Gene mean
Gene group ID, size
1, 11
2, 230
3, 3352
4, 7121
5, 3844
6, 2580
1, 26
2, 135
3, 1238
4, 6564
5, 4621
6, 4511
1, 22
2, 317
3, 3002
4, 6339
5, 3371
6, 3066
1, 29
2, 274
3, 3552
4, 6785
5, 3847
6, 3585
Scaled log−normalized expression
Gene group 1
Gene group 2
Gene group 3
Gene group 4
Gene group 5
Gene group 6

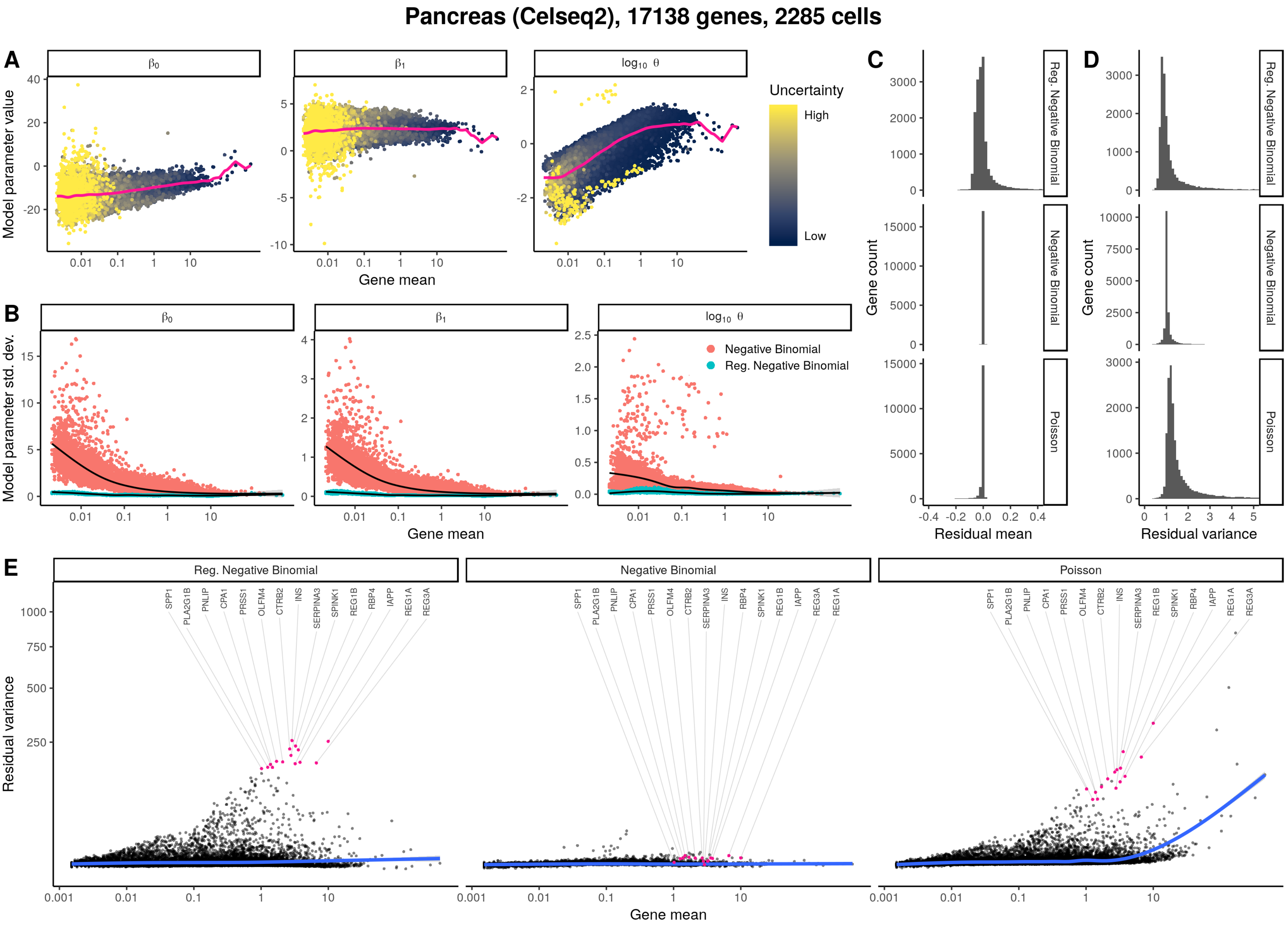

Pancreas (Celseq2), 17138 genes, 2285 cells
A
β0
β1
log10 θ
Model parameter value
Gene mean
Uncertainty
High
Low
B
Model parameter std. dev.
Negative Binomial
Reg. Negative Binomial
C
D
Reg. Negative Binomial
Negative Binomial
Poisson
Gene count
Residual mean
Residual variance
E
Residual variance
SPP1
PLA2G1B
PNLIP
CPA1
PRSS1
OLFM4
CTRB2
INS
SERPINA3
SPINK1
REG1B
RBP4
IAPP
REG1A
REG3A

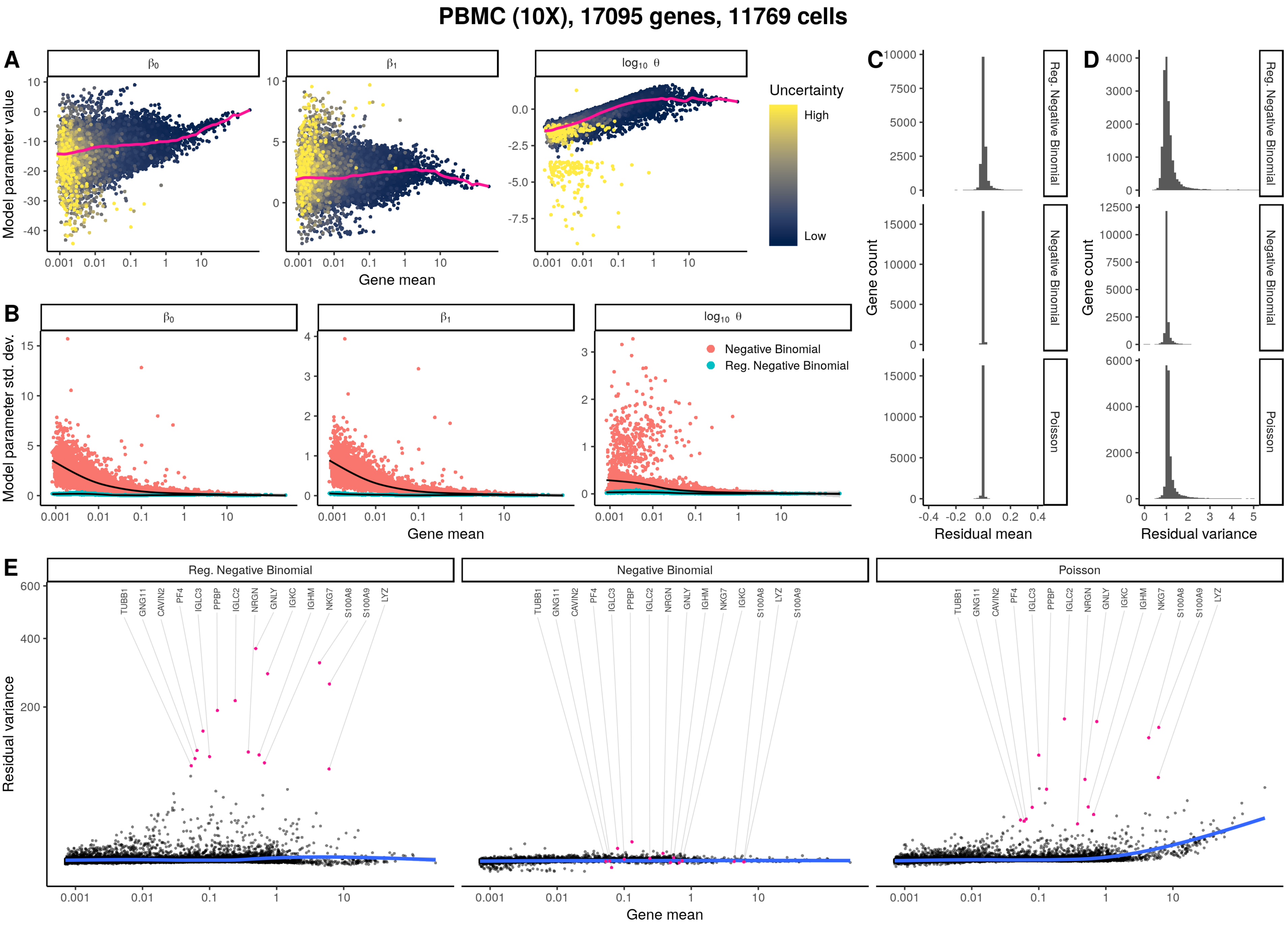

PBMC (10X), 17095 genes, 11769 cells
A
B
C
D
E
Model parameter value
Model parameter std. dev.
Gene mean
Uncertainty
High
Low
Negative Binomial
Reg. Negative Binomial
Poisson
Gene count
Residual mean
Residual variance
TUBB1
GNG11
CAVIN2
PF4
IGLC3
PPBP
IGLC2
NRGN
GNLY
IGKC
IGHM
NKG7
S100A8
S100A9
LYZ

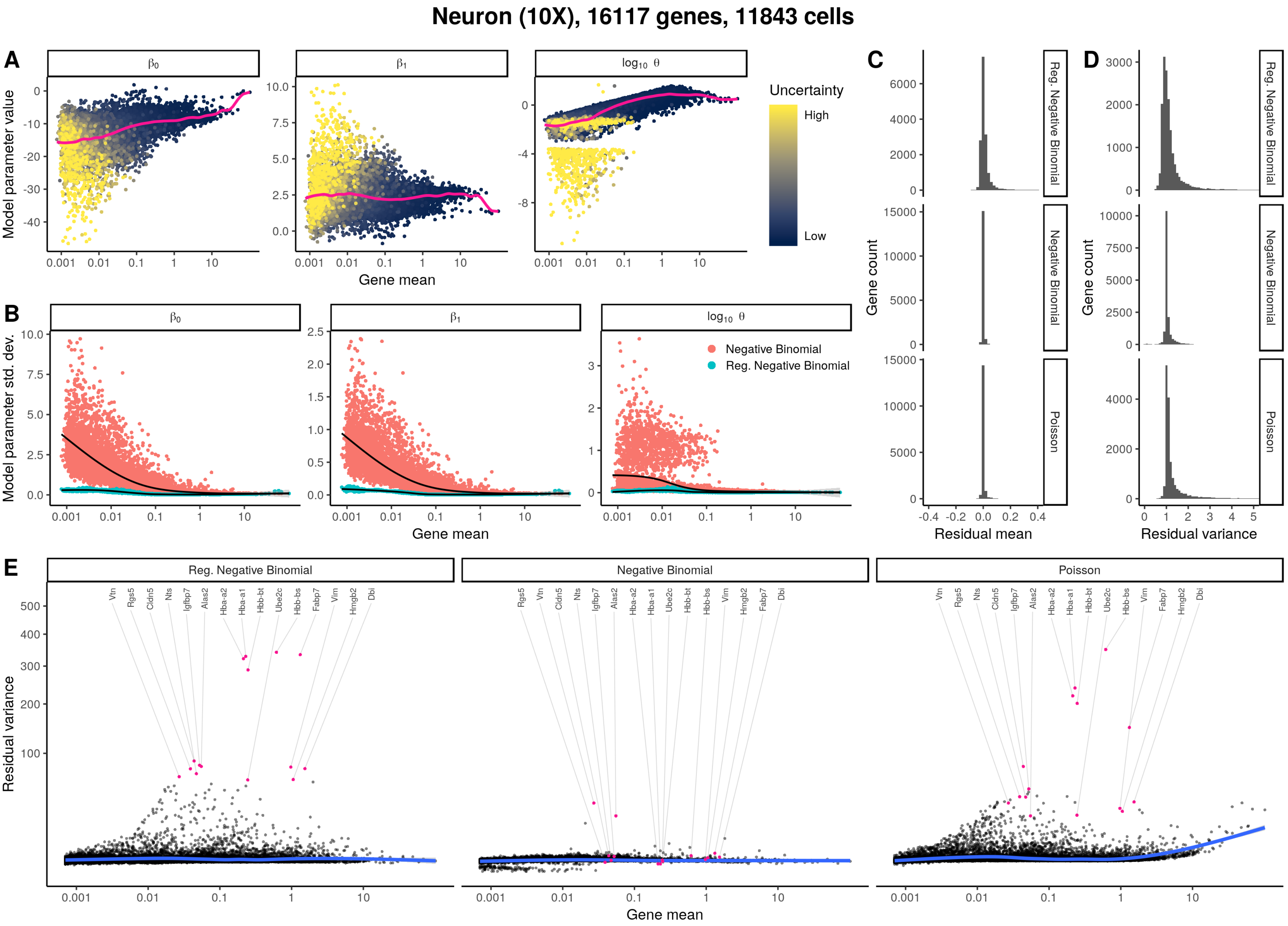

Neuron (10X), 16117 genes, 11843 cells
A
B
C
D
E
Model parameter value
Model parameter std. dev.
Gene mean
Uncertainty
High
Low
Negative Binomial
Reg. Negative Binomial
Poisson
Gene count
Residual mean
Residual variance

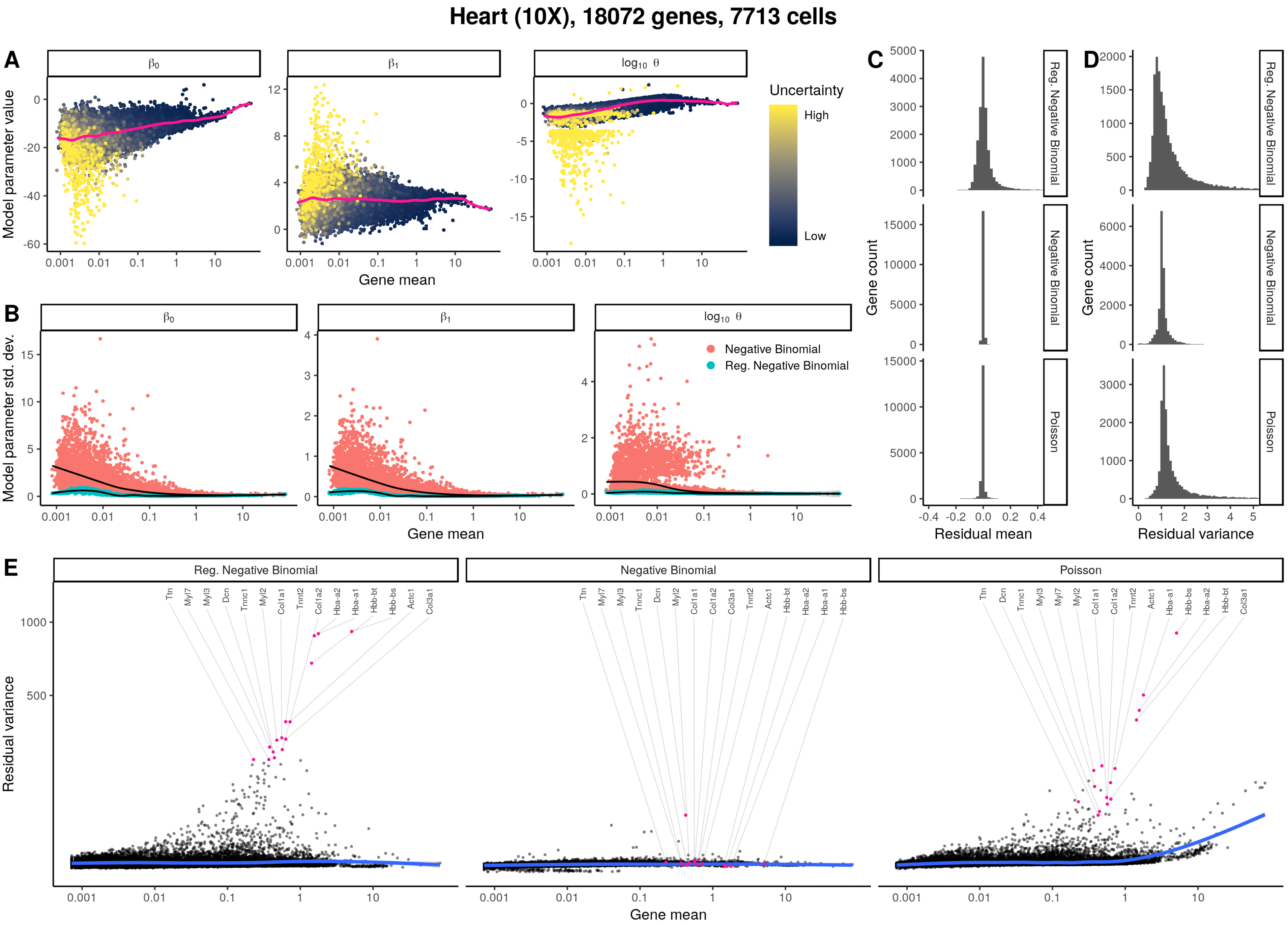
Heart (10X), 18072 genes, 7713 cells
A
B
C
D
E
Model parameter value
Gene mean
Uncertainty
High
Low
Model parameter std. dev.
Negative Binomial
Reg. Negative Binomial
Gene count
Residual mean
Residual variance
Poisson
Ttn
Myl7
Myl3
Dcn
Tnnc1
Myl2
Col1a1
Tnnt2
Col1a2
Hba-a2
Hba-a1
Hbb-bt
Hbb-bs
Actc1
Col3a1
