## Supplementary Note 2 : Using sctransform in Seurat for "Normalization and variance stabilization of single-cell RNA-seq data using regularized negative binomial regression"

*Christoph Hafemeister & Rahul Satija*

*2019-03-17*

This vignette shows how to use the sctransform wrapper in Seurat.

Install sctransform and Seurat v3

```
devtools::install_github(repo = 'ChristophH/sctransform', ref = 'develop')
devtools::install_github(repo = 'satijalab/seurat', ref = 'release/3.0')
library(Seurat)
library(sctransform)
library(ggplot2)
```

```
pbmc_data <- Read10X(data.dir = "~/Downloads/pbmc3k_filtered_gene_bc_matrices/hg19/")
pbmc <- CreateSeuratObject(counts = pbmc_data)
```

For reference, we first apply the standard Seurat workflow, with log-normalization

```
pbmc_logtransform <- pbmc
pbmc_logtransform <- NormalizeData(pbmc_logtransform, verbose = FALSE)
pbmc_logtransform <- FindVariableFeatures(pbmc_logtransform, verbose = FALSE)
pbmc_logtransform <- ScaleData(pbmc_logtransform, verbose = FALSE)
pbmc_logtransform <- RunPCA(pbmc_logtransform, verbose = FALSE)
pbmc_logtransform <- RunUMAP(pbmc_logtransform,dims = 1:20, verbose = FALSE)
```

For comparison, we now apply sctransform normalization

```
# Note that this single command replaces NormalizeData, ScaleData, and FindVariableFeatures.
# Transformed data will be available in the SCT assay, which is set as the default
# after running sctransform
pbmc <- SCTransform(object = pbmc, verbose = FALSE)
```

Perform dimensionality reduction by PCA and UMAP embedding

```
# These are now standard steps in the Seurat workflow for visualization and clustering
pbmc <- RunPCA(object = pbmc, verbose = FALSE)
pbmc <- RunUMAP(object = pbmc, dims = 1:20, verbose = FALSE)

pbmc <- FindNeighbors(object = pbmc, dims = 1:20, verbose = FALSE)
pbmc <- FindClusters(object = pbmc, verbose = FALSE)
```

Visualize the clustering results on the sctransform and log-normalized embeddings.

```
pbmc_logtransform$clusterID <- Idents(pbmc)
Idents(pbmc_logtransform) <- 'clusterID'
plot1 <- DimPlot(object = pbmc, label = TRUE) + NoLegend() + ggtitle('sctransform')
plot2 <- DimPlot(object = pbmc_logtransform, label = TRUE)
plot2 <- plot2 + NoLegend() + ggtitle('Log-normalization')
CombinePlots(list(plot1,plot2))
```

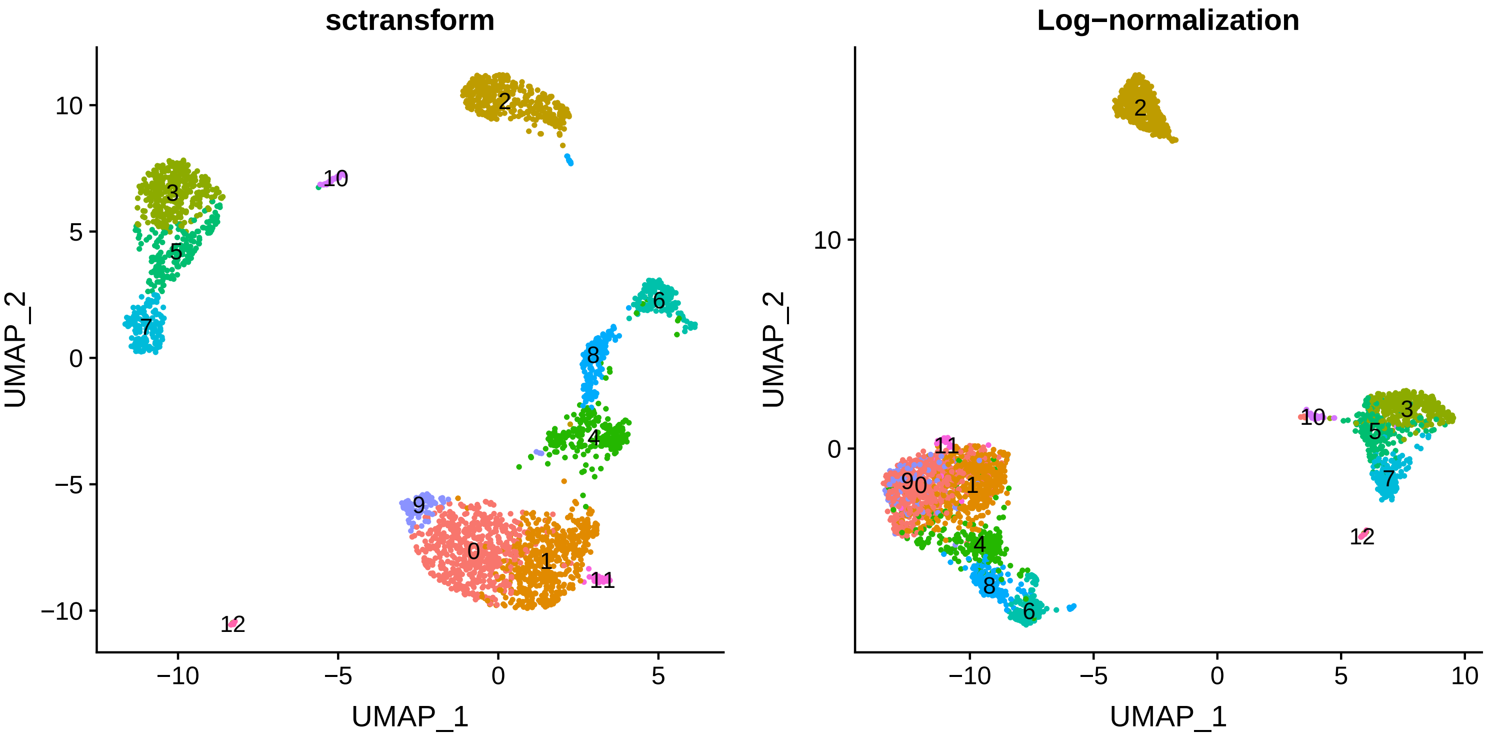


Users can individually annotate clusters based on canonical markers. However, the sctransform normalization reveals sharper biological distinctions compared to the log-normalized analysis. For example, note how clusters 0, 1, 4, 9, and 11 (all distinct clusters of T cells), are blended together in log-normalized analyses. The sctransform analysis reveals:

- Clear separation of three CD8 T cell clusters (naive, memory, effector), based on CD8A, GZMK, CCL5, GZMK expression
- Clear separation of three CD4 T cell clusters (naive, memory, IFN-activated) based on S100A4, CCR7, IL32, and ISG15
- Additional developmental sub-structure in B cell cluster, based on TCL1A, FCER2
- Additional sub-structure within NK cell cluster (CD56dim vs. bright), based on XCL1 and FCGR3A

```
# Visualize canonical marker genes as violin plots.
VlnPlot(object = pbmc, features = c("CD8A","GZMK","CCL5","S100A4","S100A4","CCR7","ISG15","CD3D"),
          pt.size = 0.2,ncol = 4)
```

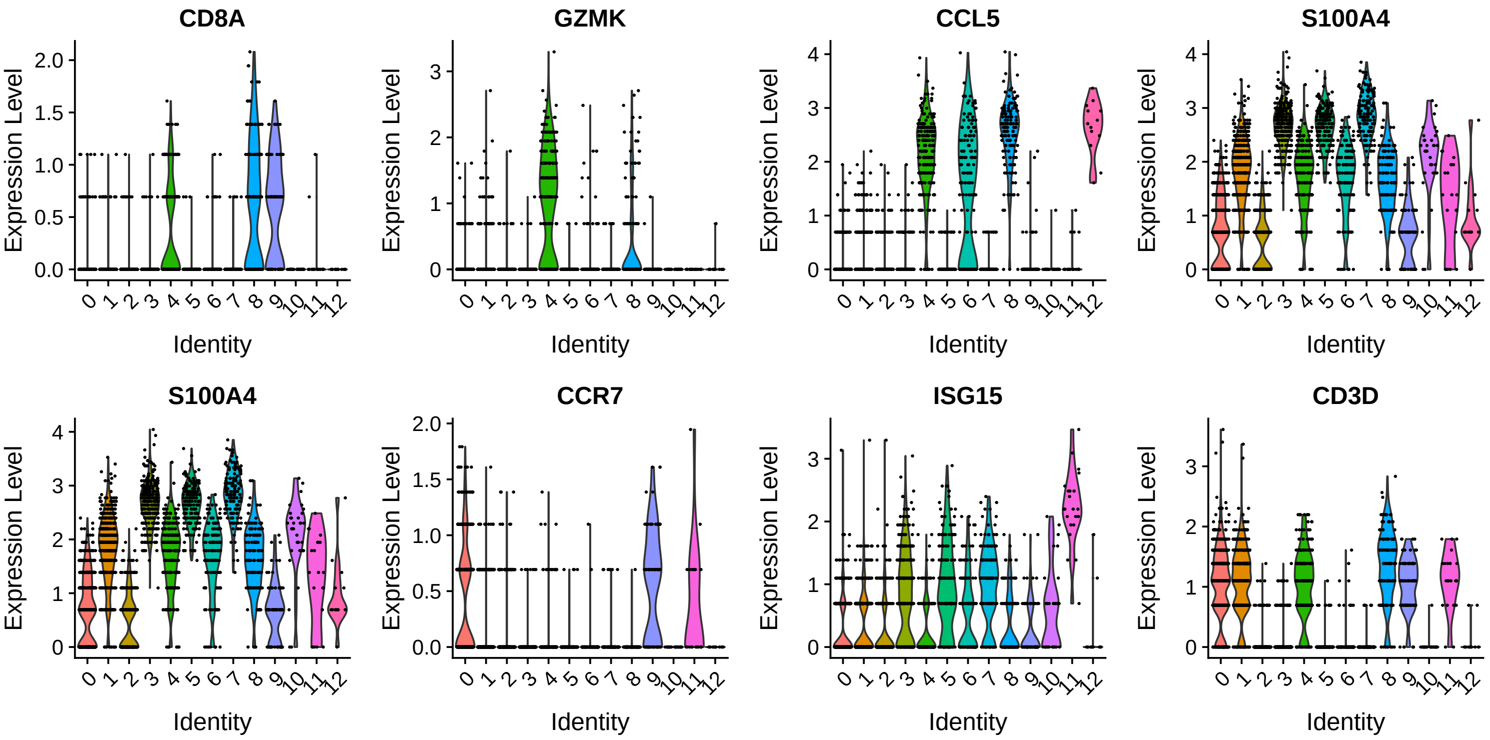

```r
# Visualize canonical marker genes on the sctransform embedding.
FeaturePlot(object = pbmc, features = c("CD8A","GZMK","CCL5","S100A4","S100A4","CCR7"),
            pt.size = 0.2,ncol = 3)
```

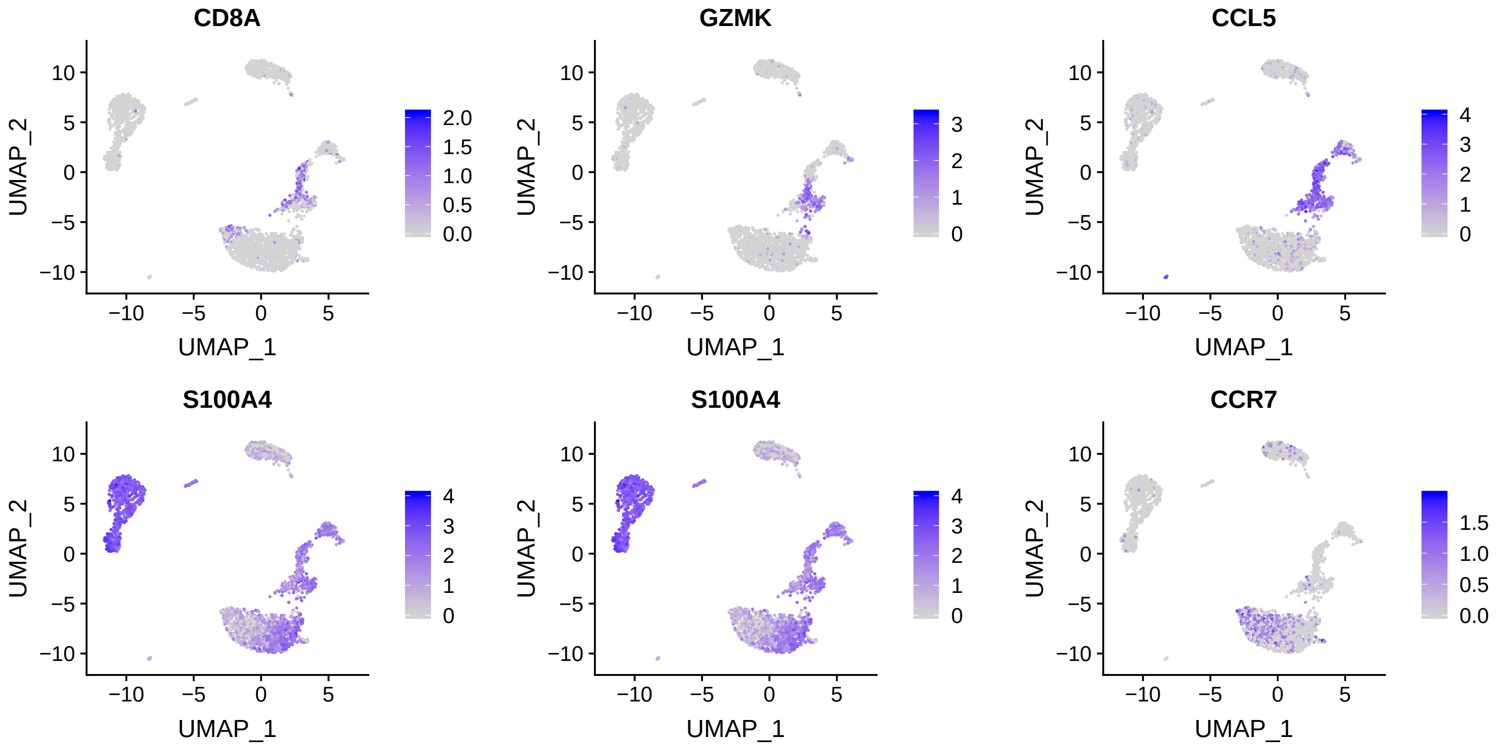


```r
FeaturePlot(object = pbmc, features = c("CD3D","ISG15","TCL1A","FCER2","XCL1","FCGR3A"),
            pt.size = 0.2,ncol = 3)
```

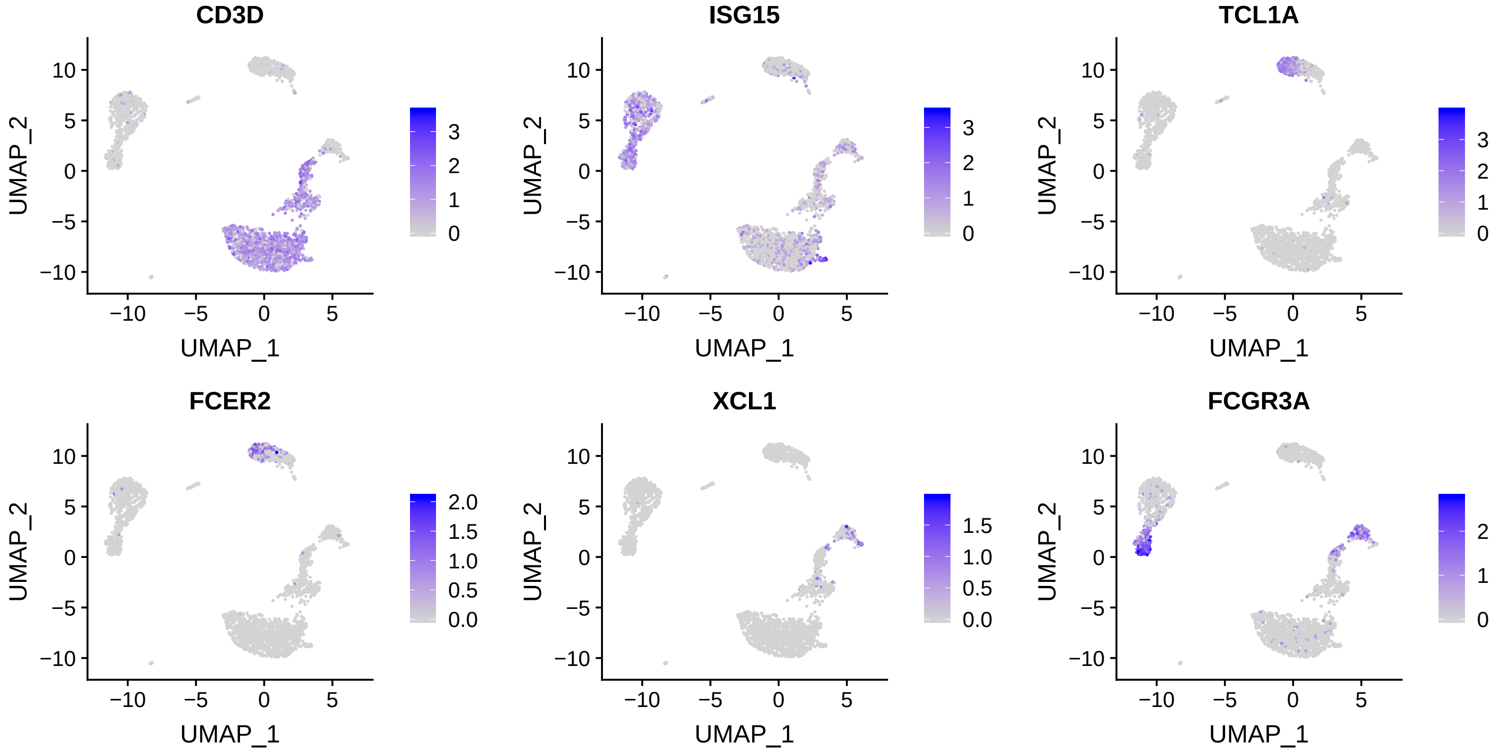
